# Stable Protein Sialylation in Physcomitrella

**DOI:** 10.1101/2020.10.09.331033

**Authors:** Lennard L. Bohlender, Juliana Parsons, Sebastian N. W. Hoernstein, Christine Rempfer, Natalia Ruiz-Molina, Timo Lorenz, Fernando Rodriguez-Jahnke, Rudolf Figl, Benjamin Fode, Friedrich Altmann, Ralf Reski, Eva L. Decker

**Author notes:** Correspondence: Eva L. Decker.

## Abstract

Recombinantly produced proteins are indispensable tools for medical applications. Since the majority of them are glycoproteins, their *N*-glycosylation profiles are major determinants for their activity, structural properties and safety. For therapeutical applications, a glycosylation pattern adapted to product and treatment requirements is advantageous. Physcomitrella (*Physcomitrium patens*, moss) is able to perform highly homogeneous complex-type *N*-glycosylation. Additionally, it has been glyco-engineered to eliminate plant-specific sugar residues by knock-out of the β1,2-xylosyltransferase and α1,3-fucosyltransferase genes (Δxt/ft). Furthermore, *P. patens* meets wide-ranging biopharmaceutical requirements such as GMP compliance, product safety, scalability and outstanding possibilities for precise genome engineering. However, all plants, in contrast to mammals, lack the capability to perform *N*-glycan sialylation. Since sialic acids are a common terminal modification on human *N*-glycans, the property to perform *N*-glycan sialylation is highly desired within the plant-based biopharmaceutical sector. In this study, we present the successful achievement of protein *N*-glycan sialylation in stably transformed *P. patens*. The sialylation ability was achieved in a Δxt/ft moss line by stable expression of six mammalian coding sequences combined with targeted organelle-specific localization of the encoded enzymes responsible for synthesis, activation, transport and transfer of sialic acid. Production of free and (CMP)-activated sialic acid was proven. The glycosidic anchor for the attachment of terminal sialic acid was generated by the introduction of a chimeric human β1,4-galactosyltransferase gene under the simultaneous knock-out of the gene encoding the endogenous β1,3-galactosyltransferase. Functional complex-type *N-*glycan sialylation was confirmed via mass spectrometric analysis of a stably co-expressed recombinant human protein.

## 1 Introduction

The biopharmaceutical sector, which in 2017 was valued at USD 1.654 billion in global sales, is continuously increasing its significant share of the global pharmaceutical market (Moorkens et al., 2017). Recombinant proteins, which represent the largest proportion of biopharmaceutical products, are manufactured only by a limited number of production platforms. These are mainly based on bacteria, yeast, insect cell, plant cell or mammalian cell cultures, whereas the latter are the most represented ones (Merlin et al., 2014). Biopharmaceuticals produced in mammalian cell cultures are in general well accepted, since these are able to perform post-translational modifications in a similar manner as in the human body (Jenkins et al., 2008).

Within mammalian systems, Chinese hamster ovary (CHO) cells are by far the most common production host (Walsh, 2018; Nguyen et al., 2020), even though they are naturally not able to link sialic acid to *N*-glycans in the human-typical predominant α2,6-manner but perform α2,3-linkages instead, which may have a destabilizing effect on recombinant antibodies (Zhang et al., 2019). Furthermore, *N*-glycans of glycoproteins produced in some non-human mammalian cell lines can carry immunogenic residues, such as the terminal *N*-glycolylneuraminic acids (Neu5Gc), a sialic acid not produced in humans (Chou et al., 2002; Tangvoranuntakul et al., 2003; Ghaderi et al., 2012) or α1,3-linked galactoses, which especially occur in SP2/0 cells (Hamadeh et al., 1992; Chung et al., 2008). Moreover, mammalian cell lines show a high heterogeneity in terms of *N*-glycosylation as well as a limited batch-to-batch reproducibility (Jefferis, 2005; Kolarich et al., 2012; Komatsu et al., 2016), which has a negative impact on the quality of a biopharmaceutical (Schiestl et al., 2011).

Alternative production platforms can offer advantages in sectors neglected by the current narrowing range of systems. In this regard, plants combine the protein processing capabilities of eukaryotic cells with cultivation requirements comparable to prokaryotic production systems, leading to lower manufacturing costs (Rozov et al., 2018). Further, plants lack human pathogens, endotoxins and oncogenic DNA sequences (Commandeur et al., 2003), and hence are generally safer than microbial or animal production hosts (Twyman et al., 2003). Between plants and animals, both the protein biosynthesis as well as the secretory pathway are highly conserved. This enables plants, in contrast to bacterial hosts, to synthesize and fold complex human proteins correctly (Tschofen et al., 2016), as well as to perform the majority of posttranslational modifications needed for high-quality biopharmaceuticals.

Due to their smaller glycome, plants display a reduced *N-*glycosylation microheterogeneity in comparison to mammalian cells (Bosch et al., 2013; Margolin et al., 2020). The latter usually synthesize a wide mixture of *N-*glycans, whereas in plants often one *N-*glycan structure predominates (Koprivova et al., 2003; Castilho et al., 2011). This leads to a rigid batch-to-batch stability and an improvement of quality and kinetics of recombinantly plant-produced biopharmaceuticals (Bosch et al., 2013; Stoger et al., 2014; Sochaj et al., 2015; Shen et al., 2016). Like mammals, plants produce complex-type *N-*glycans, which share an identical GlcNAc_2_Man_3_GlcNAc_2_ (GnGn) core but differ in some post-ER processed residues from their mammalian counterparts (Viëtor et al., 2003). The distal GlcNAc residues of mammalian *N-*glycans harbor β1,4-linked galactoses, which are often capped by a α2,6-linked sialic acid residue (Varki et al., 2015). In contrast, plant *N-*glycans are predominantly terminated with exposed distal GlcNAcs, which additionally can be decorated with an β1,3-linked galactose and an α1,4-linked fucose (Lerouge et al., 1998). This terminal structure (Galβ(1,3)(Fucα(1,4))GlcNAc), known as Lewis A (Le^a^) epitope, can be found in some plant cell-wall associated proteins (Strasser et al., 2007). It is, however, a tumor-associated carbohydrate structure in humans (Zhang et al., 2018) and it has been associated with antibody formation (Fitchette et al., 1999; Wilson et al., 2001). Additionally, the Asn-linked GlcNAc is substituted with an α1,3-attached fucose in plants, whereas in mammals this residue is α1,6-linked. Furthermore, in plants the proximal mannose harbors a bisecting β1,2-linked xylose, a sugar which is not present in mammals. Considering that the majority of biopharmaceuticals are glycoproteins and that their *N-*glycosylation plays an important role in their efficacy (Lingg et al., 2012), attention should be paid to *N-*glycosylation quality. Therefore, several model plants have been glyco-engineered in the last decades to obtain a humanized *N-*glycosylation pattern devoid of these structures (van Ree et al., 2000; Bardor et al., 2003; Gomord et al., 2005; Decker and Reski, 2012; Decker et al., 2014). Plant-specific β1,2-xylosylation and α1,3-fucosylation of *N-*glycans were eliminated in *P. paten*s by targeted knock-outs of the responsible glycosyltransferase (XT and FT) coding genes via homologous recombination (Koprivova et al., 2004). The same genes were knocked down by RNAi in *Arabidopsis thaliana* (Strasser et al., 2004), *Lemna minor* (Cox et al., 2006), *Nicotiana benthamiana* (Strasser et al., 2008), *Medicago sativa* (Sourrouille et al., 2008) and *Oryza sativa* (Shin et al., 2011). More recently, these genes were knocked out in *Nicotiana benthamiana* (Jansing et al., 2018) and *Nicotiana tabacum* BY-2 suspension cells by targeting two XT and five FT encoding genes via CRISPR/Cas9 genome editing (Hanania et al., 2017; Mercx et al., 2017). Furthermore, the Le^a^ epitope, which is a rare terminal modification of *P. patens N-*glycans, was abolished in this organism by the single knock-out of the gene encoding the responsible β1,3-galactosyltransferase 1 (GalT3; Parsons et al., 2012). Recombinant glycoproteins produced in *P. patens* with the triple KO of XT, FT and GalT3 displayed an outstanding homogeneity in the *N-*glycans, with a sharply predominant GnGn pattern (Parsons et al., 2012), providing a suitable platform for further glyco-optimization approaches. These studies also revealed a sufficient plasticity of plants towards glyco-engineering approaches without observable phenotypic impairments (Webster and Thomas, 2012).

Plant systems are already being used for the production of biopharmaceuticals. β-glucocerebrosidase, an enzyme for replacement therapy in Morbus Gaucher treatment, is obtained from carrot-cell suspensions (van Dussen et al., 2013) or ZMapp, a combination of antibodies for treatment of Ebola infections, is produced in *N. benthamiana* lacking plant-typical *N-*glycan modifications (Margolin et al., 2018). Besides these approved plant-made biopharmaceuticals there are further promising candidates in clinical trials, such as virus-like particles bearing influenza hemagglutinin proteins as *Medicago sativa*-derived vaccine against influenza (D’Aoust et al., 2010) or the moss-produced α-galactosidase for enzyme replacement therapy in Morbus Fabry treatment (Shen et al., 2016; Hennermann et al., 2019). The majority of pharmaceutically interesting glycoproteins are terminally sialylated in their native form. *N*-glycan sialylation is highly desirable due to its role in half-life, solubility, stability and receptor binding (Varki et al., 2015). However, plants are unable to perform β1,4-galactosylation, which in mammals serves as acceptor substrate for terminal *N-*glycan sialylation. Moreover, they are not able to produce, activate and link sialic acid (Zeleny et al., 2006; Bakker et al., 2008; Castilho et al., 2008), which in mammals requires the coordinated activity of six enzymes. To achieve *in planta* β1,4-galactosylation, several trials have been undertaken in different systems with varying degrees of success, including the expression of the β1,4-galactosyltransferase from humans (hGalT4) or animals or chimeric varieties (Palacpac et al., 1999; Bakker et al., 2001, 2006; Misaki et al., 2003; Huether et al., 2005; Fujiyama et al., 2007; Hesselink et al., 2014; Kittur et al., 2020; Kriechbaum et al., 2020). The efficiency and quality of galactosylation was shown to depend on the localization of the enzyme in the Golgi apparatus (Bakker et al., 2001; Strasser et al., 2009), on its expression level (Kallolimath et al., 2018) and on the investigated protein (Kriechbaum et al., 2020). *De novo* synthesis of sialic acid involves three enzymes in a four-step process within the cytosol. The biosynthesis starts with the synthesis of *N*-acetylmannosamine (ManNAc) out of its UDP-activated precursor substrate *N*-acetylglucosamine (UDP-GlcNAc), which is present in both, plants and mammals. The first reaction steps of the sialic acid production are catalyzed by the enzyme UDP-GlcNAc 2-epimerase/ManNAc kinase (GNE), which is the key enzyme of sialic acid biosynthesis (Reinke et al., 2009). GNE bifunctionally accomplishes the cleavage of UDP and the subsequent epimerization of GlcNAc to ManNAc, followed by ManNAc phosphorylation at C-6, resulting in ManNAc-6P. Subsequently, Neu5Ac 9-phosphate synthase (NANS) catalyzes the condensation with phosphoenolpyruvate (PEP) leading to Neu5Ac-9P (Roseman et al., 1961). Finally, Neu5Ac-9P is dephosphorylated by the Neu5Ac-9-P phosphatase (NANP) resulting in *N*-acetylneuraminic acid (Neu5Ac, sialic acid, Maliekal et al., 2006). Activation of Neu5Ac is then performed by the nuclear CMP-Neu5Ac synthetase (CMAS), forming CMP-Neu5Ac. This is translocated in exchange for CMP in an antiporter mechanism by the CMP-sialic acid transporter (CSAT) from the cytosol into the Golgi lumen (Aoki et al., 2003; Tiralongo et al., 2006; Zhao et al., 2006). In the final step of mammalian *N-*glycosylation, the sialyltransferase (ST) catalyzes the release of Neu5Ac from CMP and the covalent attachment of it to an β1,4-galactosylated acceptor in the (trans) Golgi apparatus, resulting in the complex-type mammalian *N-*glycosylation. In recent years several approaches have been performed to establish functional sialylation in plants. Accordingly, the simultaneous expression of the murine GNE and human NANS and CMAS coding sequences (CDS) in *A. thaliana* lead to the production of activated sialic acid (Castilho et al., 2008). In *N. benthamiana* functional *N-*glycan sialylation could be achieved in a transient approach by co-expressing five CDSs of the sialylation pathway genes, together with a chimeric GalT4 containing *N-*terminally the sub-Golgi apparatus localization determining cytoplasmic, transmembrane, and stem (CTS) region of the rat ST. This resulted in a sialylation efficiency of about 80%, detected on an additionally transiently co-expressed recombinant monoclonal antibody (Castilho et al., 2010). The first stable genetically engineered sialylating *N. benthamiana* line was described by Kallolimath et al. (2016).

Particularly, with regard to recent achievements in the production of candidate biopharmaceuticals, combined with its Good Manufacturing Practice (GMP)-compliant production capabilities, the moss *P. patens* serves as a competitive production platform for biopharmaceuticals (Decker and Reski, 2020). Preclinical trials of moss-derived recombinant human complement factor H were recently effectively accomplished (Häffner et al., 2017; Michelfelder et al., 2017), along with the moss-GAA (acid alpha-1,4-glucosidase) against Pompe disease (Hintze et al., 2020). Furthermore, the clinical phase I study of the recombinantly produced candidate biopharmaceutical moss-aGal against Fabry disease was successfully completed (Reski et al., 2018; Hennermann et al., 2019). Moss-GAA and moss-aGal proved to have better overall performance compared to their variants produced in mammalian cell cultures (Hennermann et al., 2019; Hintze et al., 2020), demonstrating the potential of *P. patens* to produce biobetters. However, the last step in humanizing *N-*glycans, protein sialylation, still has to be accomplished.

Here, we report the successful establishment of stable protein sialylation in *P. patens*, after subsequently accomplishing the synthesis of free as well as activated sialic acid, and the β1,4 linkage of the sialic acid anchor galactose to protein *N*-glycans. This will further increase the attractiveness of moss as plant-based biopharmaceutical production platform.

## 2 Materials and Methods

### 2.1 Construct generation

For cloning, coding sequences were amplified by PCR with Phusion™ High-Fidelity DNA Polymerase (Thermo Fisher Scientific, Waltham, MA, USA). All primers used in this study are compiled in Supplementary Table S1. After each PCR or restriction digest step, agarose gel purification of respective fragments was performed using QIAEX II Gel Extraction Kit (QIAGEN, Hilden, Germany) and ligations were performed using pJET 1.2 Cloning Kit or TOPO™ TA Cloning™ (Thermo Fisher Scientific), all according to the manufacturer’s protocols. Assembled vectors were verified by sequencing.

For the heterologous expression of all six genes of the sialylation pathway two multi-gene constructs, containing three PCR-amplified CDS each, were assembled via restriction-site based cloning. GNE and CSAT were amplified from murine cDNA and ST from rat cDNA. NANS, a truncated CMAS-version lacking 120 bases at the 5’ end (Castilho et al., 2010) and NANP were amplified from human cDNA. Additionally, a Blasticidin-S deaminase (BSD) expression cassette under the control of the CaMV 35S promoter and CaMV 35S terminator, provides resistance towards Blasticidin S (Kubo et al., 2013).

To assemble the multi-gene constructs, each CDS as well as the BSD cassette was amplified with restriction site-introducing primers (primers 1-14), and subcloned in the pJET 1.2 cloning vector. Expression of each glycosylation-related CDS was driven by the long CaMV 35S promoter (Horstmann et al., 2004) and the nos terminator. Promoter, multiple cloning site (MCS) and terminator were amplified via CDS-specific restriction site-introducing primers (primers 15-22) and subcloned into pJET 1.2, resulting in six different target vectors. To assemble the expression cassettes, the subcloned CDS and the multiple cloning site of the corresponding target vector were digested with the respective restriction enzymes and ligated, resulting in seven expression cassettes, containing GNE, NANS, NANP, CMAS, CSAT, ST or BSD, respectively. For generation of the two multi-gene constructs, two assembly vectors, either based on pJET 1.2 or pCR™4-TOPO® TA-vector harboring the corresponding homologous flanks for targeted genome integration and a designed MCS were created, respectively (Figures 1A and 1B). For the introduction of the first part of the sialylation pathway containing GNE, NANS and NANP (GNN) vector, homologous flanks were designed to target the integration of the construct into the *P. patens* adenine phosphoribosyltransferase (APT) gene (Pp3c8_16590V3.1). To create the multiple cloning site of the GNN-assembly vector, the APT-5’ homologous flank was amplified with a primer pair introducing an LguI restriction site at the 5’ end and AgeI, SgrDI and AvrII restriction sites at the 3’ end of the PCR product (oliogonucleotides 23 and 24). The APT-3’ homologous flank was amplified with a primer pair introducing AvrII and AscI restriction sites at the 5’ end and an LguI restriction site at the 3’ end of the PCR product (primers 25 and 26). The respective PCR-products were digested with AvrII, ligated and subsequently cloned into pJET 1.2, resulting in the GNN assembly vector (Figure 1A). GNE-, NANS- and NANP expression cassettes were subsequently introduced into the GNN-assembly vector using the AgeI + SgrDI, SgrDI + AvrII or AvrII + AscI restriction sites, respectively, resulting in the GNN-construct (Supplementary Figure S1A).

**Figure 1.**
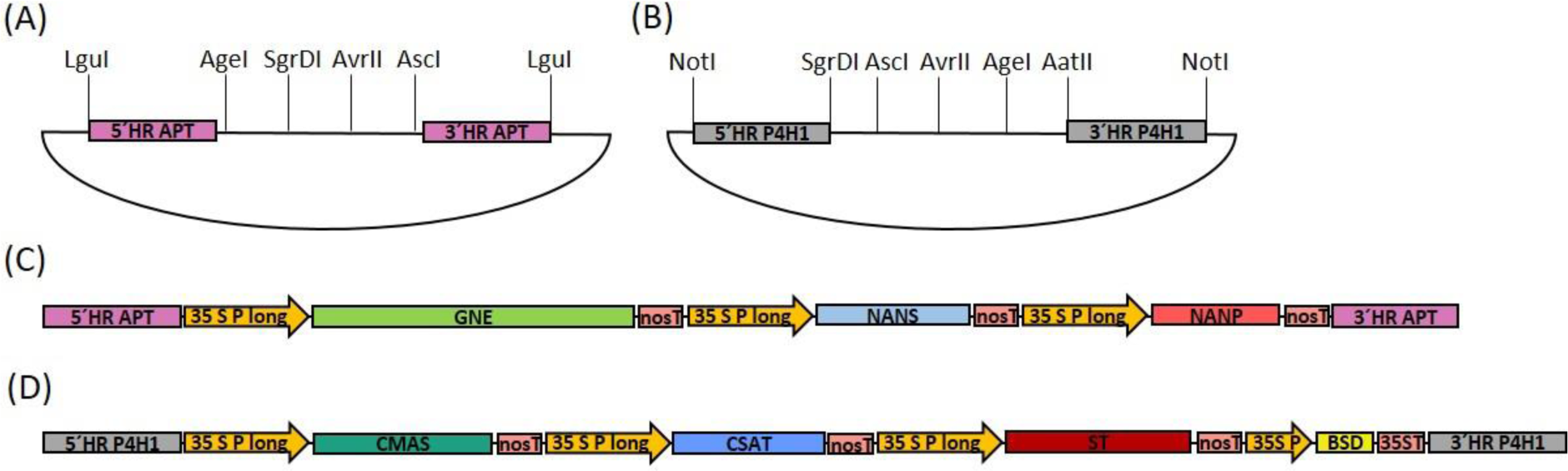
Schematic illustration of the cloned target vectors and the final GNN and CCSB constructs used for transfection. **(A)** pJET1.2-based assembly vector with a designed multiple cloning site for generation of the GNE, NANS and NANP-containing GNN construct with homologous flanks for integration into the adenine phosphoribosyltransferase (APT) locus. **(B)** pTOPO-based assembly vector with a designed multiple cloning site for generation of the CMAS, CSAT, ST and BSD-containing CCSB construct with homologous flanks for integration into the prolyl-4-hydroxylase 1 (P4H1) locus. LguI or NotI were used to linearize GNN **(C)** and CCSB **(D)** constructs, respectively, before sequential transformation of *P. patens*. 5’ HR: 5’ homologous region, 3’ HR: 3’ homologous region, 35S P long: long CaMV 35S promoter, 35S P: CaMV 35S promoter, nosT: nos terminator, 35ST: CaMV 35S terminator.

For construction of the CMAS, CSAT, ST and BSD-cassette containing multi-gene construct (CCSB) a similar strategy was used. This construct was created based on the pTOPO vector, targeted to the endogenous prolyl-4-hydroxylase (P4H1) gene (Pp3c8_7140V3.1) and additionally harbored a Blasticidin S resistance (BSD) for selection purposes. To create the multiple cloning site of the CCSB-assembly vector, the P4H1-5’ homologous flank was amplified with a primer pair introducing a NotI restriction site at the 5’ end and SgrDI, AgeI and AvrII restriction sites at the 3’ end of the PCR product (primers 27 + 28). The P4H1-3’ homologous flank was amplified with a primer pair introducing AvrII, AgeI and AatII restriction sites at the 5’ end and a NotI restriction site at the 3’ end of the PCR product (primers 29 + 30). The respective PCR-products were digested with AvrII, ligated, polyadenylated with Taq-polymerase using primers 27 and 30 and cloned into the pTOPO vector to create the CCSB-assembly vector (Figure 1B). CMAS, CAST, ST and BSD expression cassettes were subsequently introduced into the CCSB-assembly vector using the SgrDI + AscI, AscI + AvrII, AvrII + AgeI and AgeI + AatII restriction sites, respectively, resulting in the CCSB-construct (Supplementary Figure S1B). The LguI or NotI-cut GNN and CCSB constructs used for transfections are schematically illustrated in Figures 1C and 1D. A schematic overview of all GNN and CCSB cloning steps is provided in Supplementary Figure S1.

#### 2.1.1 GNE mutation

To circumvent CMP-Neu5Ac-triggered negative feedback regulation of GNE, a mutated version (GNEmut) with amino acid exchanges R263L and R266Q in the allosteric CMP-Neu5Ac binding site (Viswanathan et al., 2005; Son et al., 2011), was created. The corresponding GNE sequence alterations were performed on the previously described GNE expression construct via site-directed mutagenesis using the Phusion Site-Directed Mutagenesis Kit (Thermo Fisher Scientific) according to the manufacturer’s instructions. For complete vector amplification a mismatching primer pair, which changes the codon at position 263 from CGA to CTA and replaces the codon CGG at position 266 with CAG, was used (primers 31 + 32). After successful mutagenesis, the native GNE of the GNN-construct was replaced by GNEmut via AgeI and SgrDI restriction sites, resulting in the GM-construct.

#### 2.1.2 Cloning of a chimeric β1,4-galactosyltransferase

For cloning of a chimeric β1,4-galactosyltransferase (FTGT), the localization-determining cytoplasmic, transmembrane, and stem region (CTS) of the human β1,4-galactosyltransferase (GalT4, NM_001497.4) was replaced by 390 bp encoding the 130 amino acids comprising the CTS of *P. patens* α1,4-fucosyltransferase (FT4, Pp3c18_90V3.1), responsible for the last step in *N-*glycosylation. The CTS from the FT4 (FT-CTS) was amplified from moss cDNA with the primers 33 and 34, generating an overlapping region to the GalT4 sequence. In parallel, 927 bp comprising the catalytic domain from the GalT4 CDS were amplified with primers 35 and 36. Subsequently, the PCR products were assembled in a two-template PCR with primers 33 and 36, giving rise to the CDS of the chimeric FT-CTS:β1,4-galactosyltransferase (FTGT). For the expression of the FTGT, a bicistronic vector was cloned, where the sequence of the chimeric FTGT and the *ble* gene (Parsons et al., 2013), conferring resistance to Zeocin™ (Zeocin), were separated by the sequence of the self-cleaving 2A peptide, P2A and driven by the long CaMV 35S promoter. This sequence triggers ribosomal skipping during translation, leading to the release of two individual proteins (Donnelly et al., 2001; Kim et al., 2011). The sequence for P2A was included at the 3’ end of the chimeric FTGT CDS by two successive PCR reactions using primers 33 and 37, and 33 and 38. The CaMV 35S promoter from the Zeocin resistance cassette in the β1,3-galactosyltransferase 1-KO construct (Parsons et al., 2012) was excised with BamHI and XhoI and replaced by a long CaMV 35S promoter amplified with primers 39 and 40 using the same restriction sites. This construct bears homologous flanks targeting the endogenous β1,3-galactosyltransferase 1 locus (GalT3, Pp3c22_470V3.1) to knock it out (Supplementary Figure S2). The FTGT-P2A sequence was then inserted between promoter and *ble* gene using the XhoI site. For transfection the construct was cut using EcoRI endonuclease.

#### 2.1.3 Cloning of a chimeric α2,6-sialyltransferase

For the introduction of a chimeric sialyltransferase (FTST), consisting of the endogenous FT-CTS and the catalytic domain of the rat α2,6-sialyltransferase, a construct with homologous flanks for targeted integration into the P4H2 locus (Pp3c20_10350V3.1) (Supplementary Figure S3) was cloned via Gibson assembly. Amplification of the 5’ and 3’ homologous flanks was performed from moss genomic DNA using overhang introducing primers 41 and 42 or 43 and 44, respectively. The hygromycin selection cassette coding for a hygromycin B phosphotransferase (hpt) (Decker et al., 2015) was amplified using the primers 45 and 46. Gibson assembly was performed as described in Gibson et al. (2009) into a pJET 1.2 backbone. For introduction of the chimeric FTST variant this vector was linearized with BseRI at the 3’ end of the 5’ homologous flank and subsequently used as backbone for the introduction of the FTST anew via Gibson assembly. For this the FT-CTS together with the long CaMV 35S promoter was amplified with the primers 47 and 48 from the previously described FTGT construct and the catalytic domain of the rat ST including the nos terminator was amplified from the previously described CCSB-construct using primers 49 and 50. For transfection the construct was cut using NotI and XbaI endonucleases.

### 2.2 Plant material and cell culture

*P. patens* was cultivated as described previously (Frank et al., 2005) in Knop medium supplemented with microelements (KnopME, Horst et al. (2016)). The sialylating lines were obtained by stable transformation of the Δxt/ft moss line (IMSC no.: 40828), in which the α1,3-fucosyltransferase and the β1,2-xylosyltransferase genes have been disrupted via homologous recombination (Koprivova et al., 2004). This line additionally produces a reporter glycoprotein.

#### 2.2.1 Transfection of *P. patens* protoplasts and selection of transgenic lines

Protoplast isolation, transformation and regeneration were performed as described before (Decker et al., 2015). For each transfection event 50 µg of linearized plasmid were used. Selection for the marker-free GNN or GM lines with targeted ATP locus disruption was performed on 0.3 mM 2-fluroadenine (2-FA) containing solid KnopME plates. Protoplasts were regenerated in liquid medium for 5 days, then transferred to Knop solid medium covered with cellophane for 3 days. Subsequently, the cellophane sheets with the regenerating protoplasts were transferred onto 2-FA containing Knop plates for a further 3 weeks. Selection with Blasticidin S (Sigma Aldrich) or Zeocin (Invitrogen) were started in liquid medium on day 8 after transfection via the addition of 75 mg/L Blasticidin S or 50 mg/L Zeocin, respectively. After 4 days of selection in liquid medium the protoplasts were transferred to solid KnopME plates containing either 75 mg/L Blasticidin S or 100 mg/L Zeocin and 1% MES, respectively. The selection was done in two successive cycles of 3 weeks on selective plates interrupted by a 2-week release on non-selective KnopME plates. Hygromycin selection was performed as described before (Wiedemann et al., 2018).

### 2.3 Molecular validation of transgenic moss lines

Plants surviving the selection were screened for targeted construct integration via direct PCR (Schween et al., 2003). Successful extraction of DNA, which was performed as described before (Parsons et al., 2012), was assayed by amplifying a part of the elongation factor 1 gene (EF1α, Pp3c2_10310V3.1) with primer pair 51 and 52. Targeted integration of the GNN or GM constructs in the APT locus was confirmed using the primer pairs 53 and 54 for 5’- and 55 and 56 for 3’-integration. Targeted integration of the CCSB construct within the P4H1 locus was assayed with the primers 57 and 58 as well as 59 and 60 for the 5’- and 3’-integration, respectively. Homologous integration of the chimeric β1,4-galactosyltransferase FTGT into the GalT3 locus was confirmed using the primers 61 and 62 (5’-integration), whereas correct 3’-integration was confirmed by the primers 63 and 64. Targeted integration of the FTST construct into the P4H2 locus was assayed with the primers 65 and 66 for 5’- and 67 and 68 for 3’-integration.

### 2.4 Expression analysis of transgenic moss lines

#### 2.4.1 Real-time qPCR analysis of gene expression

To determine the expression levels of the transgenes, total RNA was isolated using Trizol (Thermo Fischer Scientific) according to the manufacturer’s instructions. RNA concentrations were determined via UV-Vis spectrometric measurement (NanoDrop ND 1000, PEQLAB Biotechnologie GmbH, Erlangen) and quality was checked via agarose gel electrophoresis. Isolated RNA was subsequently digested with DNaseI (Thermo Fisher Scientific) and cDNA synthesis was performed with random hexamers using TaqMan^®^ Reverse Transcription Reagents (Thermo Fisher Scientific), both according to the manufacturer’s protocols. Completeness of DNAseI digestion was confirmed by a non-transcribed control, without the addition of MultiScribe^®^ RT enzyme. Primer pairs were designed using the Universal Probe Library by Roche (https://lifescience.roche.com/en_de/brands/universal-probe-library.html#assay-design-center) and selected according to the lowest amount of off-target hits identified in a Phytozome (Goodstein et al., 2012) search against the *P. patens* transcriptome (V3.3; Lang et al., 2018). Oligonucleotide pair efficiencies of 2 were confirmed prior to analysis with a qPCR-run of a serial 1:2 dilution row of cDNA completed by a control without cDNA. Presence of off-targets was excluded via melting curve analysis. Measurements were prepared in white 96-multiwell plates and carried out in triplicate, using the SensiMix™ Kit and SYBRGreen (Bioline, Luckenwalde, Germany). For each triplicate cDNA amounts corresponding to 50 ng RNA were used and measurements were conducted in a LightCycler^®^ 480 (Roche) according to the manufacturer’s instructions. Absence of DNA contamination in reagents was confirmed for each primer pair with a control reaction without template addition. Amplification was performed in 40 cycles with a melting temperature of 60°C. Computational analysis of the qRT-PCR was done with LightCycler^®^ 480 software (Roche). The analysis of the expression levels of the transgenes was performed relatively against the expression values of the internal housekeeping genes coding for EF1α and for the ribosomal protein L21 (Pp3c13_2360V3.1) (Beike et al., 2015). The relative expression of the gene of interest (GOI) compared to the controls (ctrl) EF1α and L21 was calculated as 2^-ΔCT^, assuming an primer efficiency of 2, and where ΔC_T_ = C_T_GOI - C_T_ctrl and C_T_ is defined as the cycle number at which each sample reaches an arbitrary threshold (Livak and Schmittgen, 2001). Primer pairs used: GNE: 73 and 74; NANS: 75 and 76; NANP: 77 and 78; CMAS: 79 and 80; CSAT: 81 and 82; ST and FTST: 83 and 84; FTGT: 85 and 86; EF1α: 69 and 70 and L21: 71 and 72.

#### 2.4.2 RNAseq library preparation and data analysis

Total RNA was extracted from 100 mg fresh weight (FW) protonema tissue in biological triplicates for each line, using Direct-zol™ RNA MicroPrep Kit (Zymo Research, Freiburg) according to the manufacturer’s protocol. RNAseq library preparations were performed by the Genomics Unit at Instituto Gulbenkian de Ciencia (Portugal) with conditions optimized according to Baym et al. (2015), Macaulay et al. (2016) and Picelli et al. (2014) using Smart-seq2. Sequencing was performed on an Illumina NextSeq 500 instrument producing 75 bp long single-end reads.

We assessed sequence quality with FastQC (Galaxy Version 0.72+galaxy1; Available online at http://www.bioinformatics.babraham.ac.uk/projects/fastqc/ and multiQC; Galaxy Version 1.6; Ewels et al., 2016) on the public European Galaxy instance at https://usegalaxy.org (Afgan et al., 2018). For transcript quantification, reads of each library were pseudoaligned against the *P. patens* transcriptome (V3.3; Lang et al., 2018), obtained from Phytozome v12.1.5 (Goodstein et al., 2012) and complemented with the sequences of the introduced transgenes using Kallisto quant (Galaxy Version 0.43.1.4; Bray et al., 2016) for single-end reads in “unstranded” mode. According to the information obtained from the sequencing facility, the average fragment length of the libraries was specified as 340 with an estimated standard deviation of 34. For the quantification algorithm a number of 100 bootstrap samples was chosen.

### 2.5 Detection of sialic acid via periodate-resorcinol-assay

The detection of total sialic acid concentrations in moss extracts was performed according to Jourdian et al. (1971) with the following modifications: Protonema, the young filamentous tissue of *P. patens*, was harvested via vacuum filtration and frozen. Approximately 150 mg FW of frozen plant material was disrupted with a glass and a metal bead (Ø 3 mm; QIAGEN GmbH, Hilden, Germany) in a TissueLyser (MM400, Retsch GmbH, Haan, Germany) at 30 Hz for 1.5 min. For Neu5Ac extraction, the four-fold amount (v/w) of 50 mM Tris/HCl (pH 7.0) was added and the samples were vortexed for 10 min and afterwards sonicated (Bandelin Sonorex RK52, Bandelin electronic GmbH & Co. KG, Berlin, Germany) for 15 min at 4°C. Crude extracts were cleared via two subsequent centrifugation steps at 14,000 rpm and 4°C for 10 min and further 30 min centrifugation of the supernatant in a fresh reaction tube. Serial dilutions of the samples (1:2 to 1:64) and a standard row containing 1 to 40 nmol Neu5Ac (Sigma Aldrich) were prepared in extraction buffer, final volume 120 µl. For sialic acid oxidation 30 µl of a 0.032 M periodic acid solution were added, followed by 45 min incubation under gently shaking conditions at 4°C. Samples were additionally cleared via a 10 min centrifugation step at 14,000 rpm and 4°C. Subsequently, 100 µl of freshly prepared resorcinol solution (0.06% w/v resorcinol (Sigma-Aldrich), 16.8% HCl, 0.25 µM CuSO_4_) and 50 µl sample or standard were mixed in a 96-well plate (Greiner Bio-One, Frickenhausen, Germany), sealed and incubated for 45 min at 80°C. Color complexes were stabilized by the addition of 100 µl tert-butanol (Sigma-Aldrich) and the absorbance measured at 595 nm (Sunrise absorbance reader, Tecan, Männedorf, Switzerland, software Magellan™ V 7.1). Calculation of Neu5Ac concentration in the samples was performed by linear regression.

### 2.6 Detection of sialic acid (Neu5Ac) via RP-HPLC-FLD

Detection of sialic acid in moss extracts was performed as described previously for *Arabidopsis thaliana* (Castilho et al., 2008) via reverse phase high performance liquid chromatography coupled with fluorescence detection (RP-HPLC-FLD). Moss protonema tissue was dispersed with an ULTRA-TURRAX^®^ (IKA, Staufen, Germany), 1 ml of the crude extract was taken for dry weight (DW) determination and further 500 µl were mixed with 50 µl acetic acid and incubated for 10 min under shaking conditions. Extracts were cleared via 1 min of centrifugation at 16,100 x g and supernatants were further purified via a C18 SPE column (Strata C18-E, 50 mg; Phenomenex), pre-equilibrated with 1% acetic acid. After loading to the column, the samples were washed with 200 µl 1% acetic acid. The column flow-through was vacuum-dried and afterwards resuspended in 30 µl ultra-pure water. Ten µl of the C18 SPE purified extracts were DMB-labeled with 60 µl of DMB labeling reagent (Sigma-Aldrich), at 50°C for 2.5 h in the dark under shaking conditions at 750 rpm (Hara et al., 1987). Prior to analysis, quenching of the reaction was performed by the addition of 730 µl of ultra-pure water. For detection of sialic acid, 5 to 20 µl of the labelled extracts were injected on a HPLC (Nexera X2 HPLC system) with a RF-20Axs Fluorescence Detector, equipped with a semimicro flow cell (Shimadzu, Korneuburg, Austria). Separation was performed on an Aquasil C18 column (250 cm x 3 mm, 5 µm particle size; Thermo Fisher Scinetific) at a flow rate of 1 ml/min, applying a linear gradient from 30% to 42% B (70% 100 mM ammonium acetate pH 5.5, 30% acetonitrile; Solvent A was water) over 12 min and the column thermostat was set to 35°C. Fluorescence was measured with wavelengths Excitation/Emission 373 nm and 448 nm. Identification of Neu5Ac in the moss extract was performed in comparison with a DMB-Neu5Ac standard sample and confirmed by its co-eluting fluorescent profile; its concentration was determined via peak area integration.

### 2.7 Detection of activated sialic acid (CMP-Neu5Ac) via mass spectrometry

Five ml of protonema suspension culture were supplemented to an ammonia concentration of 1% (v/v) and dispersed with an ULTRA-TURRAX^®^. Afterwards the extracts were cleared via 10 min of centrifugation at 16,100 x g and supernatants were applied on a 10 mg HyperSep™ Hypercarb™ SPE cartridge (Thermo Scientific). Washing of the column was conducted with 1 ml 1% ammonia, and CMP-Neu5Ac was eluted with 600 µl 50% ACN in 1% ammonia. The elution fraction was vacuum-dried and resuspended in 10 µl 80 mM formic acid, buffered to pH 9 with ammonia. Five µl was injected on a Dionex Ultimate 3000 LC-system, using a Hypercarb™ Porous Graphitic Carbon LC Column (150×0.32 mm; Thermo Scientific). Solvent A consisted of 80 mM formic acid, buffered to pH 9, solvent B of 80% ACN in solvent A. At a flow of 6 µl/min and column oven set to 31°C, initial conditions of 1.3% B were held for 5 min, went to 19% B over 27 min and finally 63% in one minute, which was held for 7 min. The LC was directly coupled to a Bruker amaZone speed ETD ion trap instrument (Bruker, Bremen, Germany) with standard ESI source settings (capillary voltage 4.5 kV, nebulizer gas pressure 0.5 bar, drying gas 5 L/min, 200°C). Spectra were recorded in negative ion data depended acquisition mode with MS^1^ set on m/z = 613.1 which corresponds to the [M-H]^−^-ion of CMP-Neu5Ac, and simulated selected ion monitoring of m/z = 322.0 ([M-H]^−^ of CMP) was performed with MS^2^. Confirmation of CMP-Neu5Ac identity was performed via the measurement of a CMP-Neu5Ac standard demonstrating an identical fragment pattern on MS^2^ level. Quantification of CMP-Neu5Ac amounts were performed via peak area integration and compared to peak areas gained from defined amounts of a CMP-Neu5Ac standard (Sigma-Aldrich).

### 2.8 Glycoprotein analysis

#### 2.8.1 Protein extraction for mass spectrometric analyses

Six-days-old plant material (1-2 g of fresh weight), cultivated in KnopME at pH 4.5 with 2.5 mM ammonium tartrate (Sigma Aldrich) was harvested over a 100 µm sieve and resuspended in 3-fold amount extraction buffer (408 mM NaCl, 60 mM Na_2_HPO_4_x2H_2_O, 10.56 mM KH_2_PO_4_, 60 mM EDTA, 1% protease inhibitor (Sigma Aldrich), pH 7.4). The material was homogenized with an ULTRA-TURRAX^®^ for 10 min at 10,000 rpm on ice. The crude cell lysate was cleared via two successive centrifugation steps for 5 min at 5,000 rpm and 4°C. Extracts were frozen in 150 µL aliquots, in liquid nitrogen and stored at -80°C.

#### 2.8.2 Glycopeptide analysis

For mass spectrometry (MS) protein extracts were supplemented with 2% SDS and 25 mM DTT (final concentration) and incubated for 10 min at 90°C. After cooling to room temperature, proteins were S-alkylated with 60 mM iodacetamide for 20 min in darkness. Prior to SDS-PAGE the samples were mixed with 4x sample loading buffer (Bio-Rad, Munich, Germany). Separation of proteins was carried out via SDS–PAGE in 7.5% gels (Ready Gel Tris-HCl; BioRad) in TGS buffer (BioRad) at 120 V. After electrophoresis, the gel was washed three times for 10 min in water followed by a 1 h staining period with PageBlue^®^ Protein Staining Solution (Thermo Fisher Scientific). For MS analysis gel bands corresponding to high molecular weight range proteins were excised. Gel band preparations, MS measurements on a QExative Plus instrument and data analysis were performed as described previously (Top et al., 2019). Raw data were processed using Mascot Distiller V2.5.1.0 (Matrix Science, USA) and the peak lists obtained were searched with Mascot V2.6.0 against an in-house database containing all *P. patens* V3.3 protein models (Lang et al., 2018) and the sequence of the reporter protein. Glycopeptides were identified from the Mascot mgf files using a custom Perl script. Glycopeptide precursor masses were searched within these files with a precursor mass tolerance of ± 5 ppm and further validated by the presence of typical GlcNAc oxonium ions (m/z-values: [GlcNAc]^+^ = 204.087, [GlcNAc - H_2_O]^+^ = 186.076, [GlcNAc - 2H_2_O]^+^ = 168.066, [GlcNAc - C_2_H_4_O_2_]^+^ = 144.065, [GlcNAc - CH_6_O_3_]^+^ = 138.055, [GlcNAc - C_2_H_6_O_3_]^+^ = 126.055), glycan fragment ions ([GlcNAcHex]^+^ = 366.139, [GlcNAcHex_2_]^+^ = 528.191) or in the case of sialic acid containing glycopeptides the presence of Neu5Ac-specific oxonium ions (m/z-values: [Neu5Ac]^+^ = 292.103 and [Neu5Ac - H_2_O]^+^ = 274.092) (Halim et al., 2014). A fragment mass tolerance of 0.02 Da was used. A list of the glycopeptides searched and their calculated precursor masses is provided in Supplementary Table S2. Quantification of identified glycopeptides was done using a default MaxQuant search (V1.6.0.16) on the raw data. For each glycopeptide identified from Mascot mgf files, the intensity value (peak area) was extracted from the MaxQuant allPeptides.txt file by matching raw file name, precursor mass and MS^2^ scan number using another custom Perl script.

## 3 Results

### 3.1 Generation of stable sialic acid-producing lines

To establish protein sialylation in *P. patens*, six sequences encoding the enzymes required to synthesize, activate, transport and finally transfer sialic acid onto β1,4-galactosylated *N-*glycans needed to be introduced into the moss genome. These CDSs were distributed to two multi-gene cassettes, with homologous flanks for targeted integration into the *P. patens* genome (Figures 1C and 1D), of a Δxt/ft (β1,2-xylosyltransferase and the α1,3-fucosyltransferase) knock-out line expressing a reporter glycoprotein to verify the *N-*glycan sialylation of recombinant proteins.

To enable the production of sialic acid, the enzymes responsible for its biosynthesis, GNE, NANS and NANP were introduced via two versions of a multi-gene expression construct only differing in the GNE CDS used. One expression construct (GNN) contained the native GNE and in the other the GNE CDS was altered with two point mutations, R263L and R266Q, in the CMP-Neu5Ac binding site (GM). Both constructs bear homologous regions targeting the adenine phosphoribosyltransferase locus, whose disruption leads to interruption of the adenine-salvage-pathway resulting in lines resistant towards the toxic adenine analogue 2-fluoro adenine (2-FA) (Trouiller et al., 2007). The selection on 2-FA resulted in four GNN-lines with the native GNE (GNN1-4) and 64 GM-lines with the mutated GNE, which were directly screened by PCR for targeted integration of the respective construct within the genome. Homologous 5’- and 3’-construct integration was confirmed for two lines harboring the GNN-construct (GNN2 and GNN4) and 3’ homologous integration was confirmed for two further GNN-lines (GNN 1 and GNN3) as well as 26 GM-construct containing lines (GM9, 10-12, 16, 21, 22, 25-28, 36, 41, 44-47, 49, 51, 52, 57 and 61-65), respectively (Supplementary Figures S4 and S5). These lines were chosen for further experiments.

#### 3.1.1 GNN-lines produce sialic acid

In mammals, the coordinated activity of GNE, NANS and NANP leads to the synthesis of free sialic acid. To prove the activity of these enzymes in *P. patens*, sialic acid was quantified in plant extracts via a colorimetric periodate-resorcinol assay, which measures total sialic acid. Three out of four analyzed GNN-lines and 20 out of 22 analyzed GM-lines could be identified as sialic acid producers, while, as expected, the parental line does not show any signal indicating sialic acid production. Sialic acid levels between 9.8 ± 0.5 and 16.5 ± 0.7 µmol/g FW were detected in the GNN lines, whereas similar amounts ranging from 4.1 ± 0.1 up to 15.9 ± 0.5 µmol/g FW were obtained in the GM lines (Figure 2A), indicating that the epimerase and kinase activities of the enzyme with the double mutation are not affected, as previously shown in CHO cells (Son et al., 2011). For the overall best producing line GNN2 presence of sialic acid was confirmed via fluorescence detection of DMB-labeled moss extracts separated over reverse phase HPLC (RP-HPLC-FLD) compared to a DMB-Neu5Ac standard (Figure 2B). According to the peaks’ area ratio, 85 µmol Neu5Ac/g DW could be detected in GNN2 extracts, which is in agreement with the result of the colorimetric assay.

**Figure 2.**
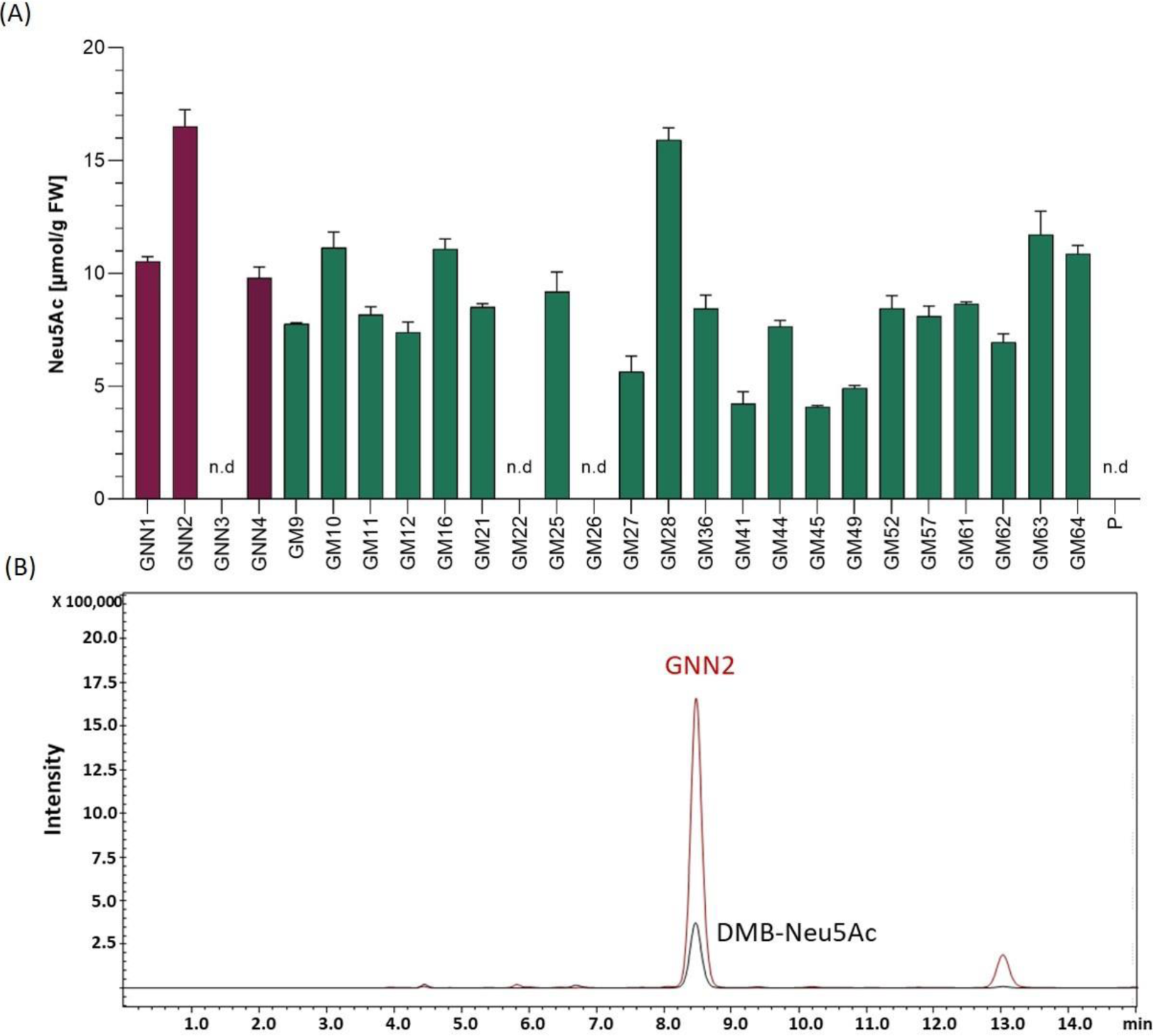
Detection of sialic acid in moss extracts of lines expressing NANS, NANP and either the native GNE (GNN lines) or the mutated version GNE_mut_ (GM lines). **(A)** Detection of sialic acid via periodate-resorcinol assay in protonema extracts of four GNN (purple), 22 GM (green) lines and the parental line (P). Neu5Ac quantification was performed in comparison to a Neu5Ac standard. Extracts were measured in duplicates in serial dilutions ranging from 1:2 to 1:32 and standard deviations (SDs) are given. Absorption was measured at 595 nm. FW: fresh weight, n.d: not detectable. **(B)** Detection of sialic acid in line GNN2 via reverse phase high performance liquid chromatography coupled with fluorescence detection (RP-HPLC-FLD). For fluorescent detection, sialic acid in the protonema extract was derivatized with 1,2-diamino-4,5-methylenedioxybenzene (DMB), resulting in DMB-Neu5Ac. Identity of the DMB-Neu5Ac in the moss extract (red line) was confirmed by comparison to DMB-Neu5Ac standard (black line).

#### 3.1.2 Expression analysis of GNE, NANS and NANP in GNN and GM-lines

Four Neu5Ac-producing lines (GNN1, GNN2, GNN4 and GM28) were characterized via Real-Time Quantitative Reverse Transcription PCR (qRT-PCR) regarding their transgene expression levels. The results are summarized in Figure 3 via the respective 2^(-ΔCT)^-values which represent the ratios between the expression level of the respective transgene compared to the expression levels of the housekeeping genes EF1α and L21. All analyzed lines strongly express the introduced transgenes, whereas the values obtained for the expression of the transgenes in the parental line are considered as background signal of the method (Figure 3). The expression level of the transgenes varied in the different lines, notably the expression level of NANP seems to be related to the amount of free sialic acid found in these lines (Figure 2A and Figure 3), indicating that the expression of this enzyme might play an important role in the efficiency of sialic acid synthesis.

**Figure 3.**
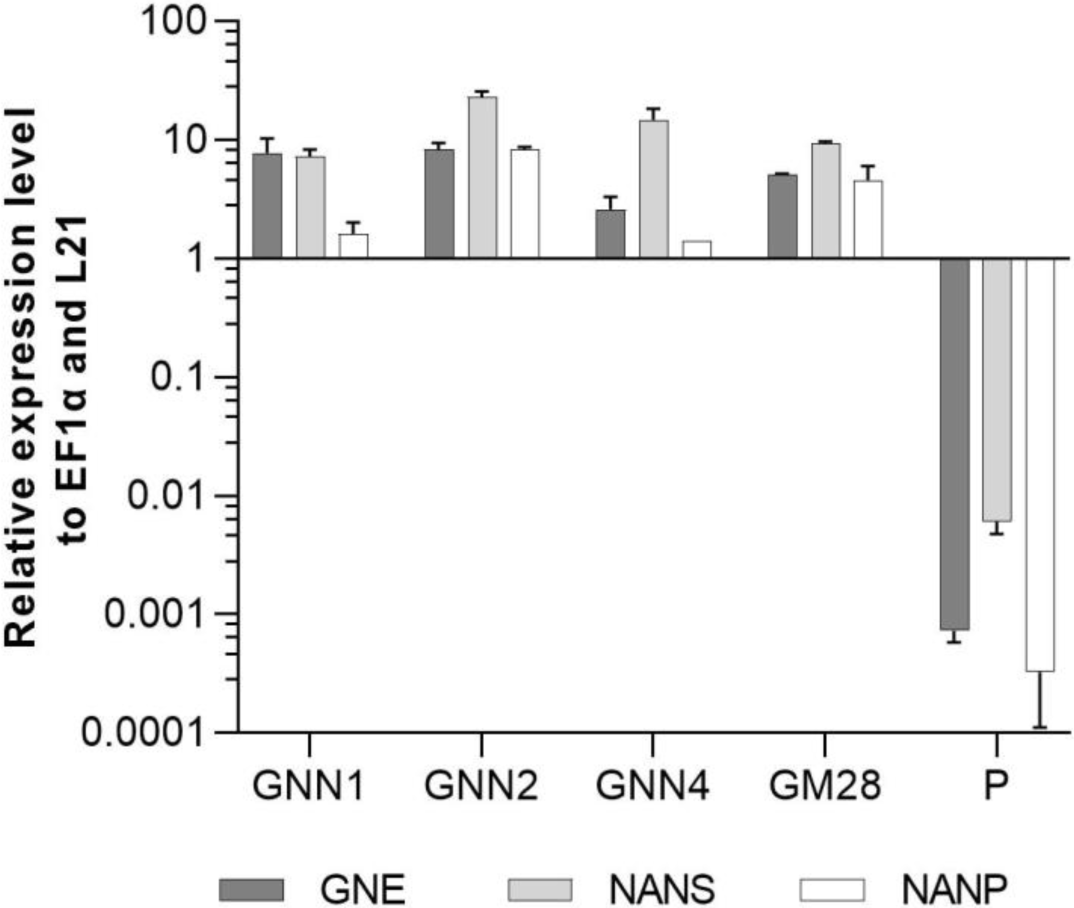
qRT-PCR based expression analysis of GNE, NANS and NANP relative to the expression of the internal housekeeping genes EF1α and L21. Expression levels of the transgenes were normalized against the expression levels of the two housekeeping genes EF1α and L21. The relative expression levels are expressed as 2^(-ΔCT)^, where ΔC_T_ = C_T_ transgene - C_T_ housekeeping genes. Bars represent mean values ± SD from n= 3. The parental plant (P) was used as negative control.

### 3.2 Generation of the complete sialylation pathway

After confirming the synthesis of sialic acid, the second part of the sialylation pathway catalyzing the activation, transport and glycosidic transfer of sialic acid was introduced. For this, the best Neu5Ac producing lines GNN2 and GM28 were transfected with the second multi-gene construct, called CCSB, containing the CDSs of CMAS, CSAT, ST as well as a Blasticidin S resistance cassette targeted to the moss P4H1 gene, to prevent plant typical prolyl hydroxylation of recombinant proteins (Parsons et al., 2013). After selection, surviving lines were directly screened by PCR for targeted integration of the CCSB-construct into the P4H1 locus. This resulted in two lines in the GNN2-background (GNC lines 4 and 7, Supplementary Figure S6) and four lines with confirmed 3’-integration in the GM28 background (GMC lines 5, 22, 23 and 46, Supplementary Figure S7).

#### 3.2.1 Expression analysis of CMAS, CSAT and ST

The expression levels of CMAS, CSAT and ST were analyzed via qRT-PCR in all seven lines with targeted integration of the CCSB-construct alongside their respective parental lines GNN2 for GNC and GM28 for GMC as negative controls. The expression levels of previously introduced GNE, NANS and NANP were analyzed anew. In four of these lines, GNC7, GMC5, GMC23 and GMC46, all six transgenes are strongly expressed, ranging from 1.4 to 100 times the expression level of the housekeeping genes EF1α and L21 (Figure 4). The other three lines showed a low expression level for at least one gene, mainly the CMAS. Unexpectedly, in GNC4 a marked decrease in the expression levels of GNE, NANS and NANP was observed.

**Figure 4.**
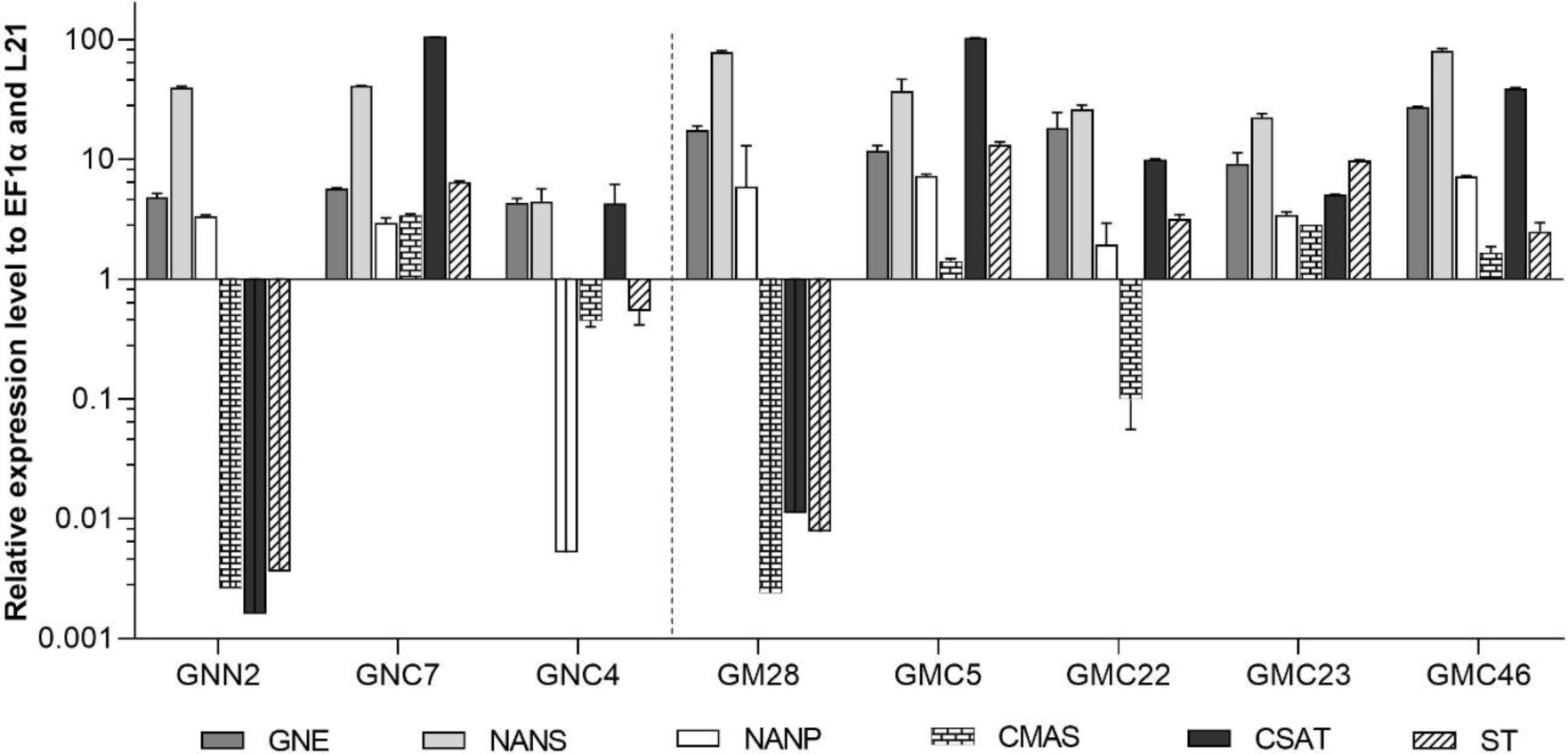
qRT-PCR based expression analysis of all six heterologous sialic acid pathway genes in two GNC lines and their parental line GNN2 and five GMC lines and their parental line GM28 relative to the expression of the internal housekeeping genes EF1α and L21. Expression levels of the transgenes were normalized against the expression levels of the two housekeeping genes EF1α and L21. The relative expression levels are expressed as 2(-ΔCT), where ΔCT = CT transgene - CT housekeeping genes. Bars represent mean values ± SD from n= 3. Lines expressing the native GNE are displayed on the left side of the dotted line, while the ones harboring the mutated GNE are depicted on the right side.

#### 3.2.2 Mass spectrometric determination of CMP-Neu5Ac in lines expressing the complete sialylation pathway

Activation of the free sialic acid by addition of CMP is a prerequisite for later transfer to the *N-*glycan structure. Therefore, the four lines robustly expressing all six transgenes (GNC7, GMC5, GMC23 and GMC46) were tested regarding their ability to activate sialic acid by measuring the CMP-Neu5Ac content via mass spectrometry in negative ion mode. The expected [CMP-Neu5Ac - H]^-^ at m/z = 613.1 on MS^1^ level and the CMP-fragment ion [CMP - H]^-^ at m/z = 322.0 on MS^2^ level could be detected in all analyzed lines, confirming the presence of CMP-Neu5Ac exemplarily shown for GMC5 in Figure 5. Verification of CMP-Neu5Ac presence in the other lines is depicted in Supplementary Figure S8. Peak area integration of the respective extracted ion chromatograms (EICs) corresponding to the m/z-value of 613.1 in comparison to the standard yielded in CMP-Neu5Ac amounts of only 2 nmol/g DW in line GNC7, in contrast 14 nmol/g DW in the lines GMC23 and GMC46 and 58 nmol/g DW in the line GMC5. This confirmed the activity of the CMAS version used in moss leading to the production of activated sialic acid. Additionally, CMP-Neu5Ac-levels in GNE mutated lines were up to 25-fold higher than those in the native GNE-harboring line GNC7.

**Figure 5.**
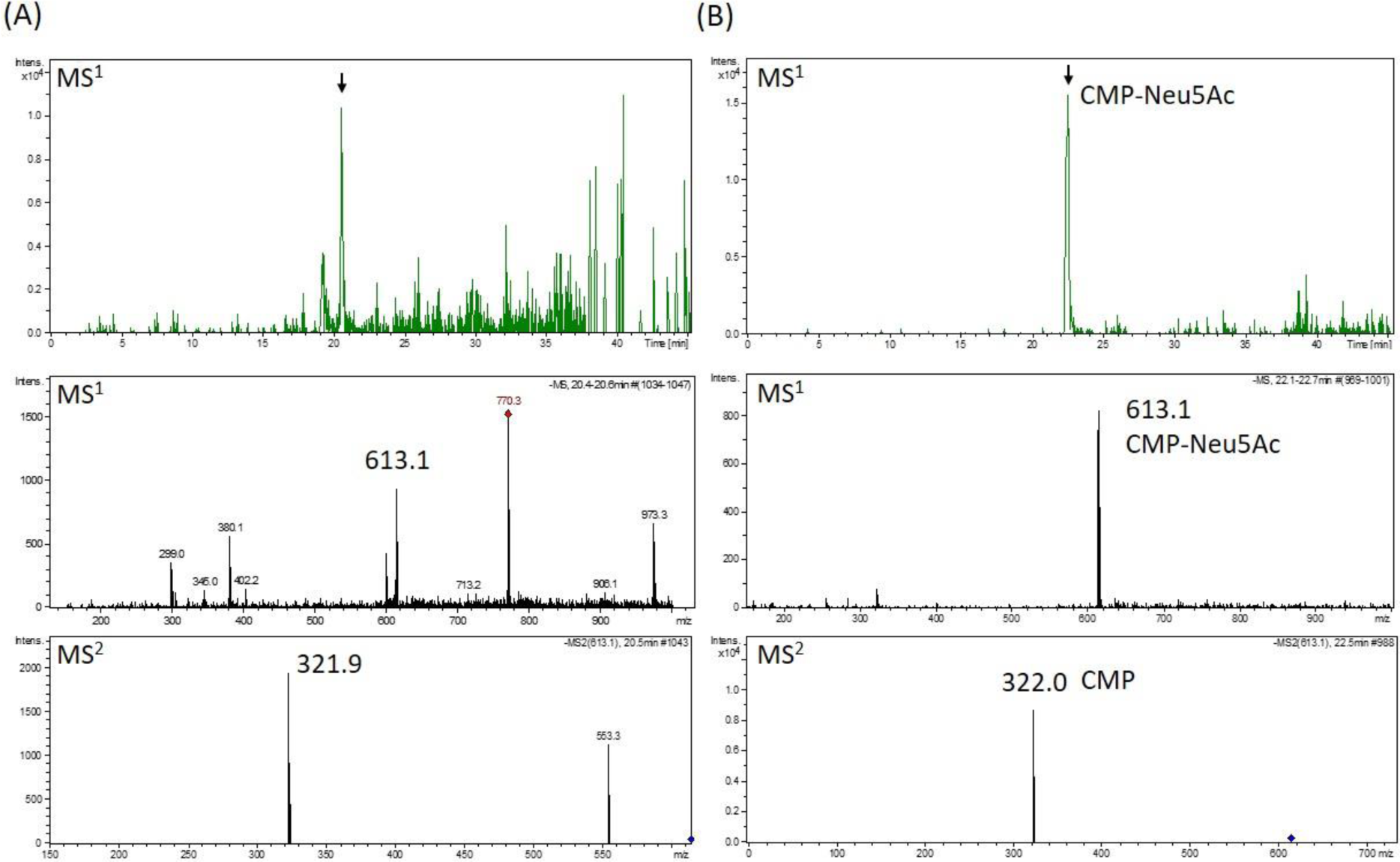
Mass spectrometric detection of CMP-Neu5Ac in a moss extract **(A)** compared to the measurement of a CMP-Neu5Ac standard (B). Extracted ion chromatogram (m/z of 613.1 ± 0.1) of LC-MS measured extract of line GMC5 (A) or 70 pmol of a CMP-Neu5Ac standard **(B)**, respectively (upper panels). In both measurements, peaks corresponding to the [M-H]^−^-ion of CMP-Neu5Ac of m/z = 613.1 could be detected on MS^1^ level (middle panels). Confirmation of corresponding MS^1^-peak identities (indicated by the black arrows) was performed on MS^2^ level via the identification of the [M-H]^−^-CMP-fragment ion of m/z = 322.0 (lower panels). CMP-Neu5Ac-concentration in GMC5 was determined via peak area integration in comparison to defined standard values and resulted in 58 nmol/g DW.

### 3.3 CMP-Neu5Ac triggers GNE feedback inhibition, which can be overcome by mutating GNE

Due to the difference in CMP-Neu5Ac amounts between plants harboring the two different versions of GNE, sialic acid content of all four selected lines expressing the six transgenes was quantified again via periodate resorcinol assay in comparison to their respective parental lines. This analysis revealed that with the introduction of the second half of the sialylation pathway the Neu5Ac content declined over 60% in the native GNE-expressing line GNC7 compared to the GNN2 parental line (Figure 6), despite no changes of the GNE, NANS and NANP expression levels being detected (Figure 4). In contrast, in the three analyzed GMC lines 5, 23 and 46, expressing the mutated GNE-version, the sialic acid content remained stable compared to the GM28 parental line (Figure 6). The higher content of CMP-Neu5Ac in lines carrying the GNE_mut_ in comparison to the native GNE containing line GNC7 indicates that the negative feedback of the activated sialic acid on the activity of GNE could be overcome with the double point mutation, which was previously described for CHO cells (Son et al., 2011). Thus, the mutated version of the enzyme is the correct choice to establish protein sialylation in moss. According to these results the GNE-mutated lines were chosen for further experiments.

**Figure 6.**
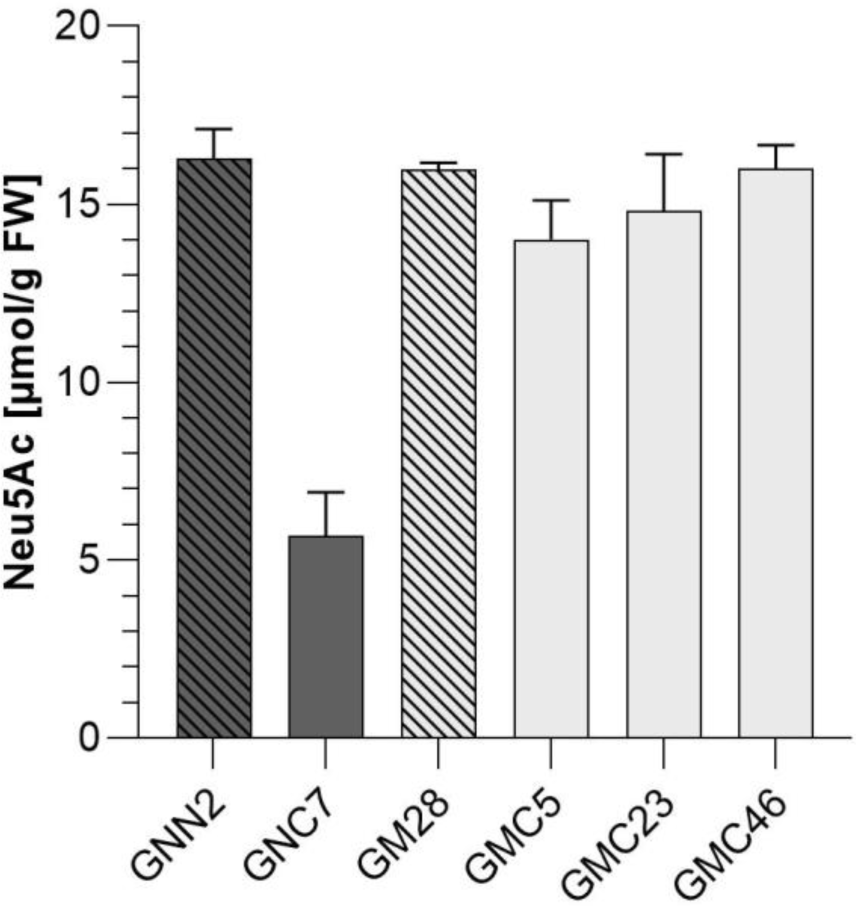
Colorimetric detection of total sialic acid content via periodate-resorcinol assay. Total sialic acid content was measured in protonema extracts of four lines expressing all six transgenes of the sialic acid pathway (GNC7, GMC5, GMC23 and GMC46, solid filled bars) and their respective parental lines GNN2 and GM28 (bars with pattern), both lacking the CMAS responsible for the activation of sialic acid. The lines expressing the native GNE are displayed in dark gray, while lines harboring the mutated GNE-version are depicted in light gray. Neu5Ac quantification was performed in comparison to a Neu5Ac standard. Extracts were measured in duplicates in serial dilutions ranging from 1:2 to 1:32 and SDs are given. Absorption was measured at 595 nm. FW: fresh weight.

### 3.4 Generation of *N-*glycan sialylation by introduction of the β1,4-galactosyl-transferase

After the introduction of the whole sialylation pathway, we aimed to provide terminal β1,4-linked galactoses for the anchoring of sialic acid to *N-*glycans. For this, a construct to simultaneously introduce the human β1,4-galactosyltransferase (GalT4) and to knock out the undesired activity of the endogenous β1,3-galactosyltransferase 1 (GalT3), responsible for the formation of Le^a^ epitopes (Parsons et al., 2012), was created. Moreover, as the glycosyltransferases in the Golgi apparatus should act in a sequential fashion, for appropriate localization of the GalT4 in the late plant Golgi apparatus (Bakker et al., 2006), a chimeric version of the enzyme was designed, bearing the CTS of the last acting enzyme in plant *N-*glycosylation, the α1,4-fucosyltransferase. This CTS region was N-terminally fused to the catalytic domain of the GalT4, resulting in the chimeric FT-CTS-GalT4 (FTGT). FTGT expression was driven by the long CaMV 35S promoter, which is according to Horstmann et al. (2004) four times stronger than the CaMV 35S promoter in *P. patens*. Line GMC23, with a high expression level of all six genes introduced previously, with the GNE_mut_ version, was transformed with the FTGT-encoding construct. After Zeocin selection, resulting lines were screened for homologous integration in the GalT3 locus, which resulted in four lines with targeted FTGT-construct integration (GMC_GT19, GMC_GT25, GMC_GT28 and GMC_GT80, Supplementary Figure S9). These four lines as well as the line GM28 as a negative control, were analyzed for the respective expression level via qRT-PCR. All lines expressed the chimeric β1,4-galactosyltransferase strongly, between 29 and 155 times higher than the housekeeping genes EF1α and L21 (Figure 7).

**Figure 7.**
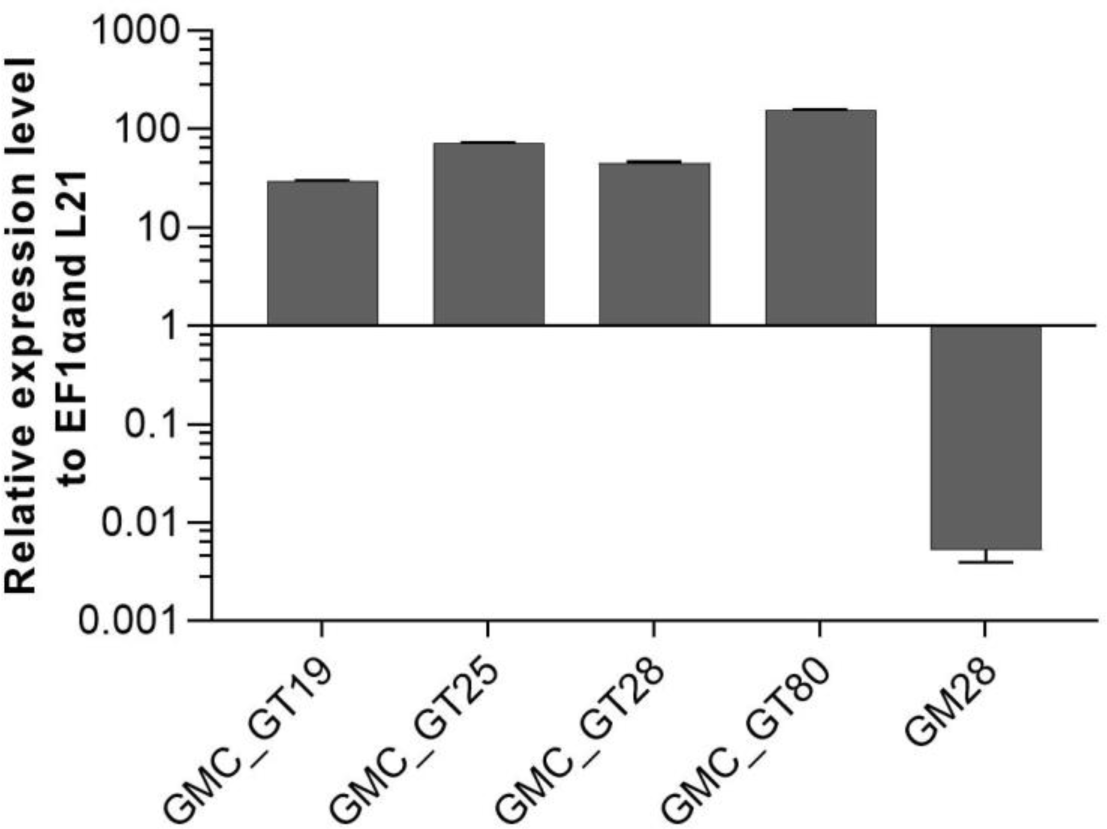
qRT-PCR based expression analysis of chimeric FTGT relative to the expression of the internal housekeeping genes EF1α and L21. Expression levels of the chimeric β1,4-galactosyltransferase (FTGT) were determined with a primer pair targeting the catalytic domain of the human GalT4 and normalized line internally against the expression levels of the two housekeeping genes EF1α and L21. The relative expression levels are expressed as 2^(-ΔCT)^, where ΔCT = CT transgene - CT housekeeping genes. Bars represent mean values ± SD from n= 3.

### 3.5 Mass spectrometric analysis of the *N-*glycosylation pattern of complete sialylation lines

The efficiency of galactosylation and subsequent sialylation of *N-*glycans in the FTGT-expressing lines GMC_GT19, 25, 28 and 80, was assessed on glycopeptides of the stably co-expressed reporter protein via mass spectrometry. Evaluation of the *N-*glycosylation pattern was performed via integration of MS^2^-confirmed glycopeptide elution profiles on MS^1^ level. Our present glycopeptide analysis does not allow us to distinguish conformational isomers of *N*-glycans but enables us to make conclusions about their composition. For simplification in the description of the results, we present all possible isomers of an *N*-glycan combined under one structure, e.g. AM, MA or a mixture of both, are all displayed together as AM. The MS-analyses revealed that the investigated lines displayed a galactosylation efficiency between 45% and 60% within the analyzed glycopeptides (Figure 8). Among galactosylated glycopeptides, 80% corresponded to AM-structures. On average, 8% of the *N-*glycans were biantennary galactosylated (indicated by AA, Figure 8). Further, our data indicate that the degree of bi-galactosylated peptides is higher in plants with lower expression level of the FTGT (Figure 7 and Figure 8). On glycopeptides harboring galactoses mass shifts of 132.0423 Da or 264.0846 Da were observed in up to 50% of all detected galactosylated *N-*glycans. This suggests the linkage of one or two pentoses to the newly introduced β1,4-linked galactoses (Figure 8 and Supplementary Figure S10). Non-galactosylated glycopeptides carried almost exclusively the complex-type GnGn glycoform. *N-*glycans bearing plant specific xylose, α1,3-linked fucose, Le^a^ epitopes or the corresponding aglycons were not detected for any of the analyzed glycopeptides. Despite the fact that with the introduction of a β1,4-galactosyltransferase into a CMP-Neu5Ac-producing and CSAT and ST-expressing moss line all requirements for sialylation are addressed, in none of the MS-analyzed lines sialylation of *N-*glycans was detectable. A corresponding result was reported for *N. benthaminana* by Kittur et al. (2020).

**Figure 8.**
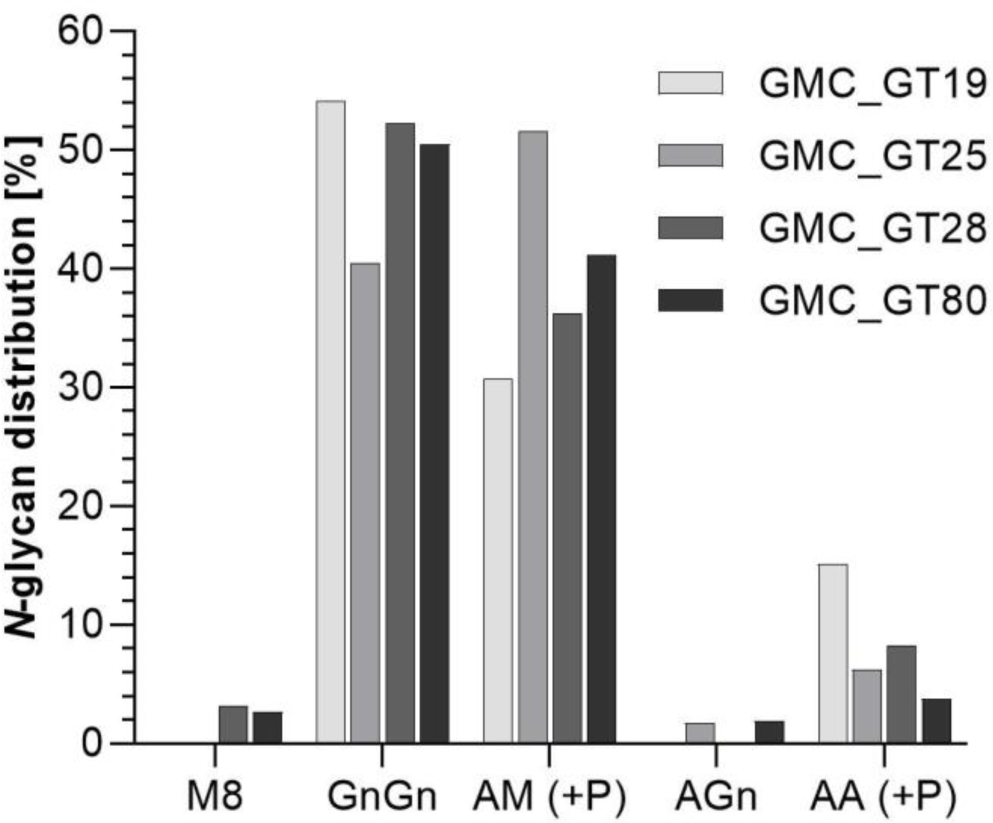
Mass spectrometric analysis of glycosylation patterns in GMC_GT lines. MS-based relative quantification of glycopeptides identified on the tryptically digested reporter glycoprotein of complete sialyation lines. Relative quantification was based on peak area integration of extracted ion chromatograms (EICs) on MS^1^ level, for which peak identities were confirmed on MS^2^ level. For quantification, areas of all confirmed peaks per measurement were summed up and the relative percentages are given for each identified glycan structure. M: mannose, Gn: *N-*acetylglucosamine, A: galactose, (+P): indicates that a variable proportion of the corresponding *N-*glycan is decorated with pentoses.

### 3.6 RNAseq-analysis revealed the drop of ST expression after FTGT introduction

To address the question why no sialic acid was attached to *N-*glycans in the GMC_GT lines harboring all genes necessary for sialylation, we analyzed the changes in gene expression in the line GMC_GT25, and its parental line GMC23, without the chimeric β1,4-galactosyltransferase. RNAseq analysis revealed that the introduction of FTGT led to a decrease of CMAS and CSAT expression of approximately 95% and nearly abolished the ST expression, whereas the expression levels of the first three pathway CDSs remained stable (Figure 9). Even if the reason for the striking drop in ST expression level could not be explained, we assume that this lack of expression is responsible for the absence of sialylation in the GMC_GT lines.

**Figure 9.**
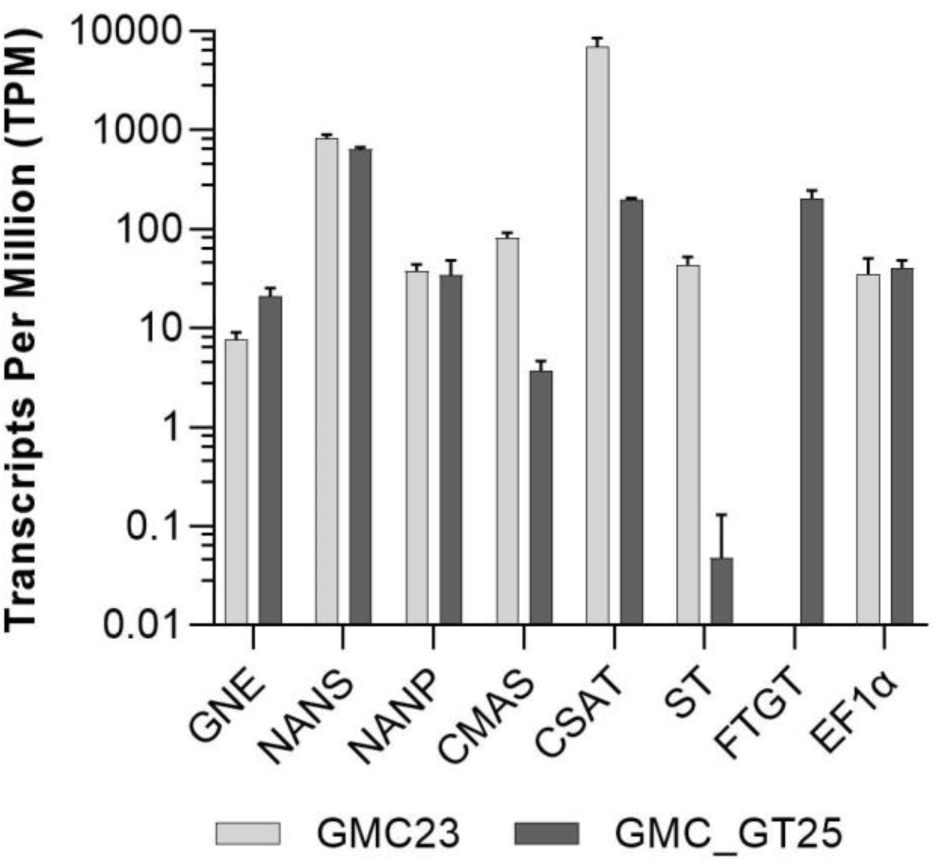
RNAseq expression analysis. Expression levels of all seven pathway related heterologous sequences and the strongly expressed housekeeping gene EF1α in the lines GMC23 and GMC_GT25. Bars represent mean values ± SD from n = 3biological triplicates.

### 3.7 Lines strongly expressing the FTST are able to perform stable *N*-glycan sialylation

To overcome the absence of ST expression in GMC_GT25, the line was transformed with a chimeric ST version. In this variant, the original CTS of the rat ST was replaced by the CTS of the endogenous α1,4-fucosyltransferase already used for the FTGT, giving raise to the chimeric variant FTST. The integration of the FTST was targeted to the P4H2 locus, which is part of the plant-prolylhydroxylases family (Parsons et al., 2013). Plants surviving the hygromycin-based selection were analyzed for targeted construct integration within the P4H2 locus, which could be confirmed for eight GMC_GT_FTST lines (GMC_GT_FTST2, 12, 13, 24, 64, 69, 78 and 79; Supplementary Figure S11). Although these plants express several transgenes and display multiple modifications, no obvious impairment in growth could be assessed.

FTST expression was quantified via qRT-PCR in all GMC_GT_FTST lines with targeted genome integration. The used primers were targeted to the catalytic domain of the sialyltransferase, thus amplifying both ST variants. Therefore, the lines GMC_GT_FTST, carrying both the native and the chimeric ST were compared to their parental line GMC_GT25, only expressing the native ST version. As previously detected, the GMC_GT25 displayed only a very weak ST expression, while six out of the eight tested GMC_GT_FTST lines strongly express the chimeric sialyltransferase, in a range between 23 and 104 times the expression level of the housekeeping genes EF1α and L21 (Figure 10).

**Figure 10.**
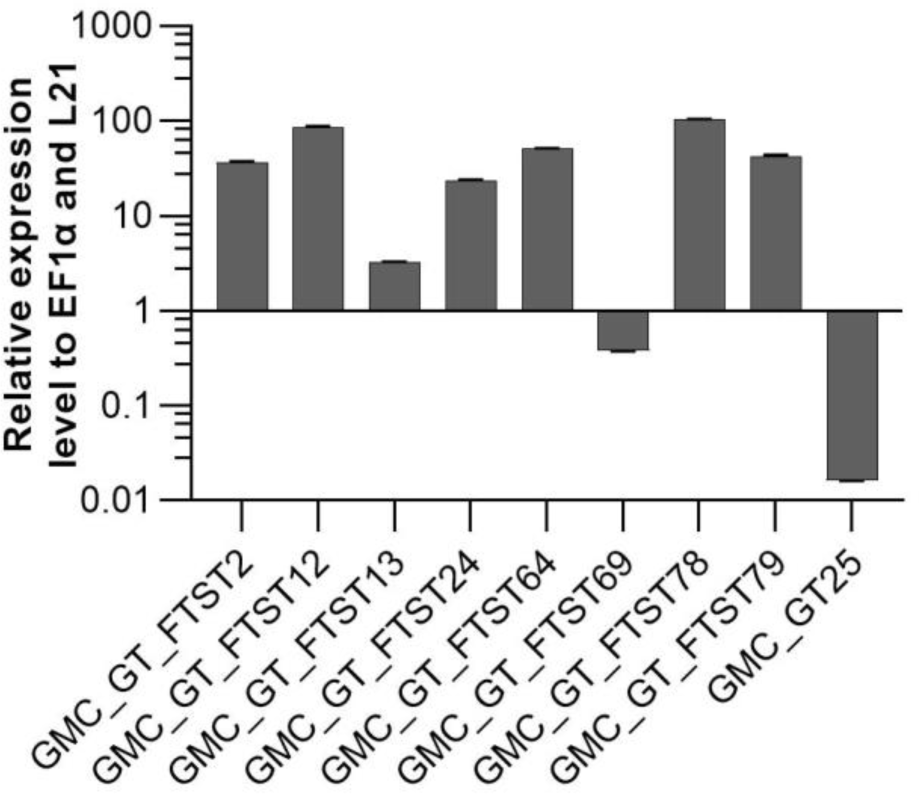
qRT-PCR based expression analysis of chimeric FTST variant relative to the expression of the internal housekeeping genes EF1α and L21. Expression level of the FTGT variant was determined with a primer pair targeting the catalytic domain of the rat ST. FTST-expression levels were normalized line internally against the expression levels of the two housekeeping genes EF1α and L21. The relative expression levels are expressed as 2^(-ΔCT)^, where ΔCT = CT transgene - CT housekeeping genes. Bars represent mean values ± SD from n= 3.

To investigate FTST enzyme activity, three lines with FTST expression levels ranging from low to high (GMC_GT_FTST13, 64 and 78) were chosen for MS-based glycopeptide analysis. This analysis revealed that the galactosylation efficiency increased from 60% in the GMC_GT25 parental line to 85-89% in FTST expressing lines (Supplementary Figure S12). Moreover, a higher share of biantennary galactosylated *N-*glycans could be detected in all analyzed lines after the introduction of the FTST, increasing from 6.3% in GMC23_GT25 to e.g. 33.7% in GMC_GT_FTST64 (Supplementary Figure S12). Further, in the two moderate and strong FTST expressing lines GMC_GT_FTST64 and 78 stable *N-*glycan sialylation could be confirmed, whereas in the weak-expressing line GMC_GT_FTST13 no *N-*glycan sialylation was detectable (Figure 11A and Supplementary Figure S12). Verification of *N*-glycan sialylation was performed via the detection of defined glycopeptide m/z-ratios on MS^1^ level (Supplementary Figures S13-S16) as well as the detection of sialic acid reporter ions [Neu5Ac - H_2_O]^+^ with m/z=274.092 and [Neu5Ac]^+^ with m/z=292.103 and fragments of the corresponding peptide backbone on MS^2^ level (Figure 11B-11D and Supplementary Figures S13-S16). In GMC_GT_FTST78, the line with the highest FTST expression level, sialylation could be detected in all three analyzed glycopeptides (Figure 11B-11D). Altogether, sialic acid was linked to 6.3% of all detected glycopeptides, of which 6.0% were NaM, while the remaining 0.3% displayed NaGn structures. Further, 70.8% AM and 0.8% AGn structures were detected, but almost half of them were decorated with one or two pentoses. Interestingly, no pentoses were found when these *N*-glycans were sialylated. Biantennary galactosylation was present in 11.3% of the glycopeptides, but no sialylation of this structure could be detected. GMC_GT_FTST64, the analyzed line with the intermediate FTST-expression level, displayed an overall sialylation efficiency of 1% NaM structures (Supplementary Figure S12), indicating that the efficiency of sialylation is dependent on the expression level of the sialyltransferase. These results verified the correct synthesis, localization and sequential activity of seven heterologous mammalian enzymes and confirmed that stable *N-*glycan sialylation is possible in moss.

**Figure 11.**
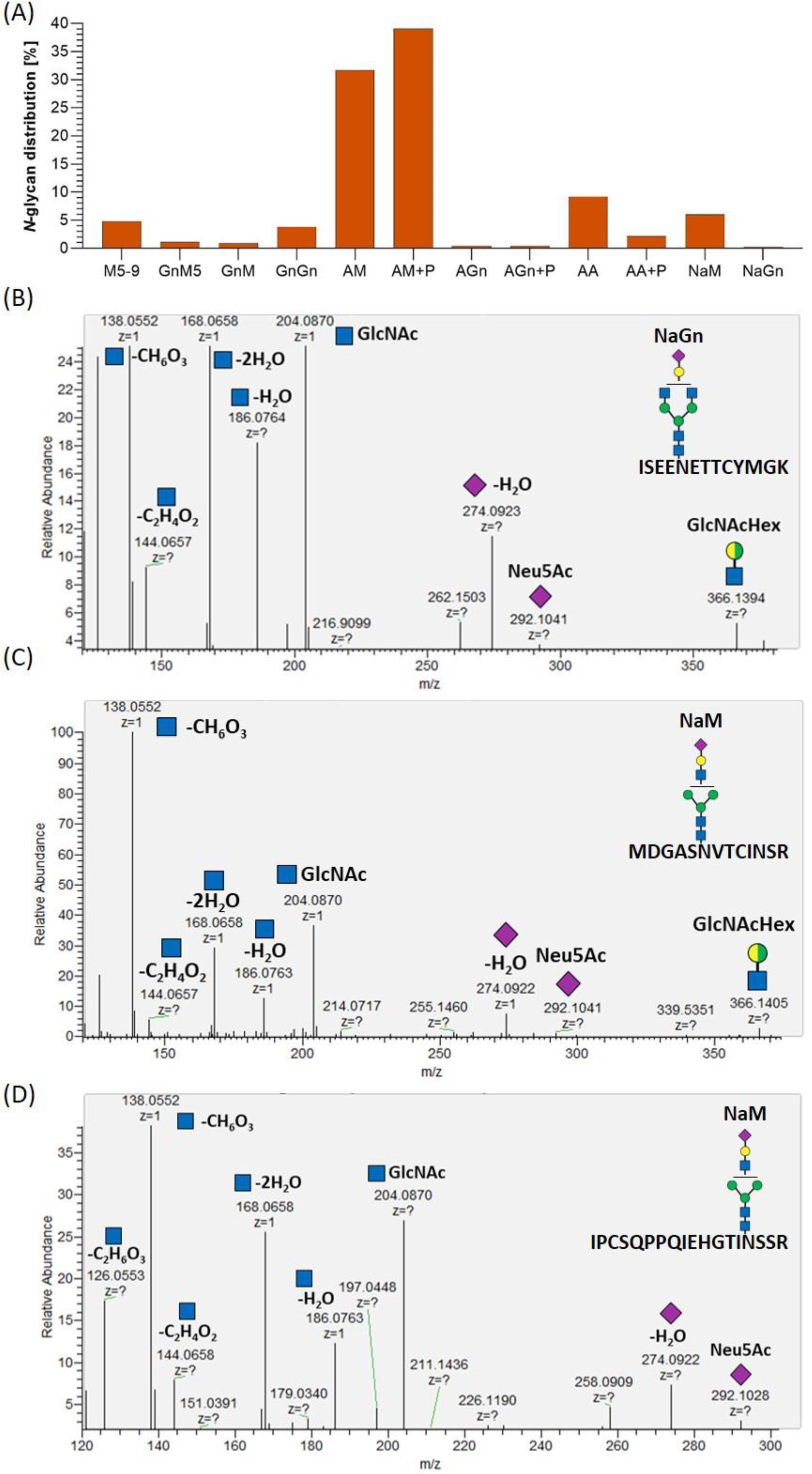
Mass spectrometric analysis of the glycosylation pattern in GMC_GT_FTST78 and MS^2^-based verification of sialylated glycopeptides. **(A)** MS-based relative quantification of glycopeptides identified on the tryptic digested reporter glycoprotein of the complete sialyation line GMC_GT_FTST78 with a chimeric sialytransferase. Relative quantification was based on peak area integration of extracted ion chromatograms (EICs) on MS^1^ level, for which glycopeptide identities were confirmed on MS^2^ level. For quantification, areas of all confirmed peaks per measurement were summed up and the relative percentages are given for each identified glycan structure. **(B-D)** MS^2^-based verification of *N*-glycan sialylation on all three analyzed tryptic glycopeptides by the identification of *N-*acetylglucosamine (GlcNAc) and sialic acid (Neu5Ac) reporter ions with the following m/z-values: [GlcNAc]^+^ = 204.087, [GlcNAc - H_2_O]^+^ = 186.076, [GlcNAc - 2H_2_O]^+^ = 168.066, [GlcNAc - C_2_H_4_O_2_]^+^ = 144.065, [GlcNAc - CH_6_O_3_]^+^ = 138.055, [GlcNAc - C_2_H_6_O_3_]^+^ = 126.055), [Neu5Ac]^+^ = 292.103, [Neu5Ac - H_2_O]^+^ = 274.092 and the detection of the glycan fragment ion [GlcNAcHex]^+^ = 366.139.M: mannose (green circle), Gn: *N-*acetylglucosamine (blue square), A: galactose (yellow circle), Na: sialic acid (purple rhombos), Hex : Hexose (yellow and green circle: stands for the presence of either mannose or galactose).

## 4 Discussion

Since many therapeutics are sialylated glycoproteins, the ability to perform sialylation on complex-type *N-*glycans is highly required for biopharmaceutical production platforms. *N-*glycan sialylation plays an important role for the plasma half-life of glycoproteins, as it protects them from clearance (reviewed in Varki, 2008), which is highly favorable to achieve long lasting therapeutic effects.

To achieve *N-*glycan sialylation in *P. patens* several challenges need to be addressed, namely the production of free sialic acid (Neu5Ac) from the precursor UDP-*N-*acetylglucosamine, which is also present in plants (Stanley et al., 2015), its activation to CMP-Neu5Ac, its transport to the Golgi apparatus and finally its transfer onto a β1,4-galactosylated *N-*glycan. This requires the coordinated activity of seven mammalian enzymes localized in three different plant subcellular compartments including the nucleus, cytosol and Golgi apparatus. Stable integration of the respective genes was directed via gene targeting, which is a frequently used tool for genome engineering in *P. patens* (e.g. Khraiwesh et al., 2010; Sakakibara et al., 2013; Decker et al., 2015; Horst et al., 2016; Wiedemann et al., 2018). These knock-in approaches were aimed at selection or to avoid undesired plant-typical posttranslational modifications. The adenine phosphoribosyltransferase is an enzyme involved in the purine salvage pathway and recycles adenine into AMP, but can also recognize adenine analogues as substrates, which are toxic when metabolized (Schaff, 1994). Knock-out of the single coding gene for this enzyme in *P. patens* results in resistance to adenine analogues as 2-fluoro adenine; an effect that has been employed as a marker for homologous recombination frequency (Trouiller et al., 2007; Kamisugi et al., 2012). We used this strategy to positively select GNN-transformed plants avoiding the introduction of an additional foreign gene for selection. The integration of the CCSB- and FTST-expression cassettes was targeted to the genes coding for the prolyl-4-hydroxylase 1 (P4H1) and the prolyl-4-hydroxylase 2 (P4H2), respectively, which are members of the plant P4H-family, responsible for hydroxylation of prolines in consensus sequences different to those recognized in humans (Parsons et al., 2013). Moreover, as hydroxyproline serves as anchor for plant-typical O-glycosylation, the formation of these structures should be avoided in biopharmaceuticals. Besides, the generation of the potential immunogenic Le^a^-epitope was abolished in *P. patens* by targeting the FTGT-expression cassette to the gene encoding β1,3-galactosyltransferase 1 (Parsons et al., 2013).

The production of free sialic acid was achieved by the simultaneous expression of the sequences encoding the first three cytosolic active enzymes of the sialic acid biosynthesis pathway: murine GNE, human NANS and human NANP, resulting in the production of Neu5Ac in 10 times higher amounts than previously reported for plants (Castilho et al., 2008). In *A. thaliana* and *N. benthamiana* transformed with GNE and NANS, endogenous activity of a putative plant NANP was sufficient for the synthesis of Neu5Ac (Castilho et al., 2008). However, our data indicates that in moss additional NANP-expression levels lead to higher Neu5Ac production.

As the full-length CMAS-CDS could not be amplified from human cDNA, a shorter version, where 120 bp at the 5’ end are missing (Castilho et al., 2008), was cloned and introduced in sialic acid-producing lines. Moss plants expressing the truncated version of CMAS were able to activate sialic acid, confirming its activity in *P. patens*.

The activity of GNE, the first enzyme in the biosynthesis of sialic acid, is known to be inhibited by the CMAS product CMP-Neu5Ac (Kornfeld et al., 1964). Mutated versions of this enzyme (R263L and/or R266Q /R266W) were already used in *N. benthamiana*, insect and CHO cells to overcome this negative feedback and increase sialylation of recombinant proteins (Viswanathan et al., 2005; Bork et al., 2007; Son et al., 2011; Kallolimath et al., 2016). We expressed in moss the GNE_mut_ version carrying the double mutation R263L, R266Q which was previously described to be more active *in vitro* than the single mutated versions (Son et al., 2011). Sialic acid-synthesizing moss plants carrying GNE_mut_ produced similar amounts of sialic acid as plants expressing the native GNE-version, confirming that the R263L, R266Q mutations in the allosteric site of GNE do not affect the epimerase/kinase activities of the enzyme. In contrast to the native GNE-expressing lines, no decrease in the production of sialic acid was observed in GNE_mut_ expressing lines after the introduction of CMAS, revealing that the negative feed-back loop of GNE was successfully prevented in moss. Consequently, up to 25 times more CMP-Neu5Ac was achieved in moss plants expressing the GNE_mut_ compared to lines with the native GNE.

The 2A peptide sequences derived from the *Picornaviridae* family are known to cause a ribosomal skipping event during the process of translation (Szymczak et al., 2004), resulting in isolated proteins from a single open reading frame. The ability of *P. patens* to recognize the 2A peptide sequence from the foot-and-mouth disease virus has already been reported (Pan et al., 2015). We used the P2A sequence (Kim et al., 2011), originated from the porcine teschovirus-1, another virus belonging to this family, to separate the transgene of interest, in our case the chimeric β 1,4-galactosyltransferase, from the *ble* resistance gene. This resulted in the synthesis of independent FTGT and the Bleomycin resistance protein, verified by their corresponding enzymatic activities.

Different qualities and degrees of galactosylation have been reported so far after the introduction of the β1,4-galactosyltransferase in plants. Enzymes involved in *N-*glycosylation are localized in the ER and/or in the Golgi apparatus according to their sequential manner of action in the *N-*glycan maturation, and this localization is determined by their N-terminal CTS region (Saint-Jore-Dupas et al., 2006). Early incorporation of galactose into the maturing *N-*glycan before the action of mannosidase II leads to hybrid-type glycans (Schachter et al., 1983; Hesselink et al., 2014). Therefore, we aimed to target the β1,4-galactosyltransferase activity to the trans Golgi apparatus by creating a chimeric enzyme consisting of the catalytic domain of the human β1,4-galactosyltransferase and the localization-determining CTS region of the moss endogenous α1,4-fucosyltransferase (FT4), which is the last enzyme known to act on the plant *N-*glycans maturation (Saint-Jore-Dupas et al., 2006). Lines expressing this synthetic GalT4 variant (FTGT) displayed up to 60% of galactosylation on the reporter glycopeptides. Mono- and biantennary galactosylated glycans were detected, up to 50% and 15%, respectively. In addition, in some of the galactosylated *N-*glycans the appearance of mass shifts corresponding to the molecular weight of one or two pentoses occurred, indicating that β1,4-galactosylated *N-*glycans were decorated with additional sugars of thus far unknown identity. The characterization of these residues remains a future task. A very similar galactosylation pattern, including the appearance of pentoses, was described by Kittur et al. (2020) on erythropoetin in transgenic *N. tabacum* lines expressing a chimeric GalT4 with the CTS region of the rat α2,6-sialyltransferase (STGT). Interestingly, in transgenic Δxt/ft *N. benthamiana* lines the same STGT variant led to predominantly bi-galactosylated *N-*glycan structures on a transiently co-expressed antibody (Strasser et al., 2009). However, the extent of biantennary galactosylation also seems to depend on the investigated glycoprotein, as in contrast to the high galactosylation efficiencies obtained on antibodies, poor levels were achieved for other proteins in the same system (Kriechbaum et al., 2020). Further, our results indicate that the expression level of the enzyme plays a role in the quality of galactosylation. Earlier studies performed in *N. benthamiana* indicated that there is an optimal GalT expression level and levels beyond it resulted in higher amounts of immature *N-*glycans (Kallolimath et al., 2018). This is in agreement with our results, since the amount of bi-galactosylated *N-*glycans was higher in lines with a lower FTGT expression level. Therefore, an improvement of the glycosylation pattern towards a higher share of bi-galactosylated *N-*glycan structures via modulation of promoter strength or number of inserted copies in stably transformed moss plants is a future task.

Achieving stable sialyation in *P. patens* requires not just the correct subcellular localization and coordinated activity of seven mammalian enzymes involved in this process, but also their optimal expression levels to produce fully processed glycans with terminal sialic acid. Since the same promoter was used to express all genes, the differences in the expression levels observed between moss transgenic lines should mainly be determined by the number of constructs integrated into the genome or the loci of integration, respectively. The use of homologous flanks to target the integration in *P. patens* do not always limit this recombination process to the intended locus alone. In contrast to transient systems, the process of plant selection according to the expression level of the gene of interest is a necessary step. According to RNAseq data, the introduction of the FTGT resulted additionally in a drop of CMAS and CSAT and a dramatically decrease of ST expression levels. It has been reported that during moss transfection rearrangements may appear, such as small deletions, insertions or concatenation of the constructs, which are characteristic of non-homologous end-joining or homologous recombination pathways (Kamisugi et al., 2006; Murén et al., 2009). These events might be an explanation for the marked decrease in the expression level of these genes, since some copies of the transgenes might be damaged by recombination events. However, we currently cannot explain why ST expression was especially affected.

To circumvent the loss of ST expression, we created and inserted the chimeric gene for FTST into the genome of the production line. The efficient expression of this chimeric ST variant led to stable sialylation of the reporter glycoprotein. The observed structures were NaM and NaGn *N-*glycans, or their isomers, appearing on all analyzed glycopeptides. Furthermore, FTST expression led to an increased galactosylation efficiency from 60 up to 89% combined with an increase in the share of biantennary galactosylated *N-*glycans. Although the sialylation levels achieved in this study are still low (around 6.3%), this is an important step towards the production of sialylated, high-quality biopharmaceuticals in *P. patens*. Different strategies need to be addressed to further optimize the expression levels of the genes involved in the sialylation pathway. It is also well known that culture conditions affect the glycosylation pattern in CHO cells, such as culture pH, temperature, media and supplements, operation mode of the bioreactor and feeding strategies (Hossler et al., 2009; Lewis et al., 2016; Wang et al., 2018; Ehret et al., 2019). Therefore, evaluating the influence of cell culture factors might be a further strategy to increase sialylation levels in *P. patens*.

We established stable *N-*glycan sialylation in *P. patens* by the introduction of seven mammalian enzymes, some of them as chimeric versions, resulting in a sialylated co-expressed human glycoprotein. This demonstrates the coordinated activity and correct subcellular localization of the corresponding enzymes within the cytosol, the nucleus and the Golgi apparatus. Stable expression of all transgenes involved in both the production of the reporter protein as well as the glycosylation, enables the complete characterization of the production line and ensures consistent product quality. The now established ability of stably transformed *P. patens* to sialylate proteins widens the range of *N-*glycosylation patterns attainable in this system considerably. This facilitates the customized production of recombinant biopharmaceuticals with various degrees of post-translational modifications to ensure maximal activity. This makes moss a versatile and attractive production system within the biopharmaceutical industry.

## 5 Author Contributions

LLB performed most of the experiments, SNWH performed the MS data analysis, CR performed the RNAseq data analysis, TL cloned the GNN, GM and CCSB constructs and FRJ cloned the FTGT construct, RF performed the mass-spectrometric based Neu5Ac and CMP-Neu5Ac analyses, LLB, JP, NRM, FA, BF, RR and ELD designed the study and wrote the manuscript.

## 6 Funding

This work was supported by the Excellence Initiative and Strategy of the German Federal and State Governments (GSC-4 SGBM to FRJ and EXC 294 BIOSS to RR), Eleva GmbH (to RR) and the Deutscher Akademischer Austauschdienst DAAD (to NRM).

## 7 Acknowledgements

We thank Agnes Novakovic, Ingrid Heger, Kai Clemens Carsten Gaa and Maxime Karrer for their support to this work, Prof. Dr. Mitsuyasu Hasebe for kindly providing the BSD plasmid, Dr. Karsten Häffner for providing human and rat cDNA, Dr. Asifa Akhtar for providing murine cDNA, and the Genomics Unit of Instituto Gulbenkian de Ciencia for RNAseq analyses. We thank Prof. Dr. Bettina Warscheid for the possibility to use the QExactive Plus instrument and Anne Katrin Prowse for proof-reading of the article.

## 8 Conflict of Interests

The authors are inventors of patents and patent applications related to the production of recombinant proteins in moss. RR is an inventor of the moss bioreactor and a founder of Greenovation Biotech, now Eleva GmbH. He currently serves as advisory board member of this company.

## Supplementary Material

**Supplementary Table S1.**
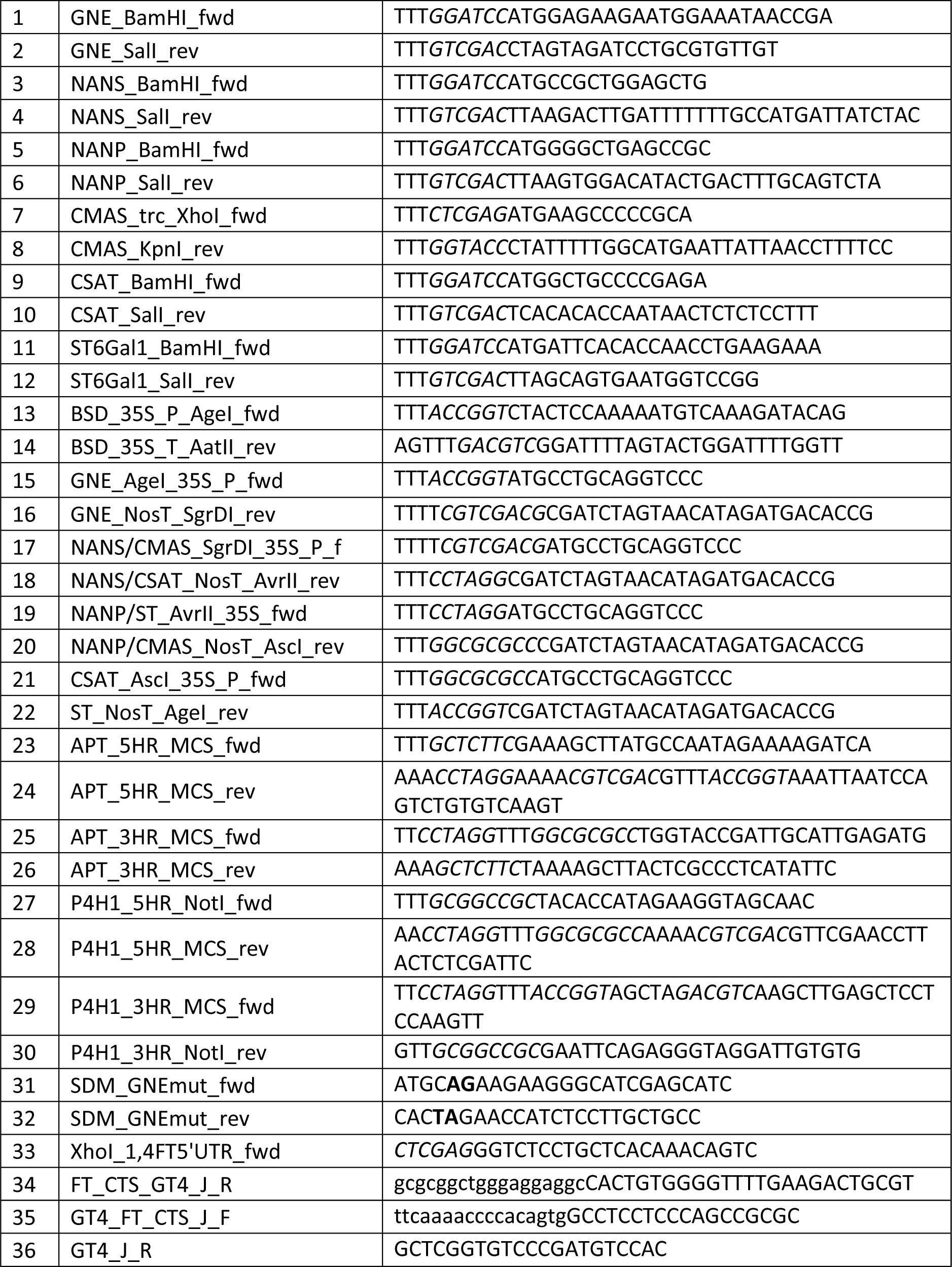
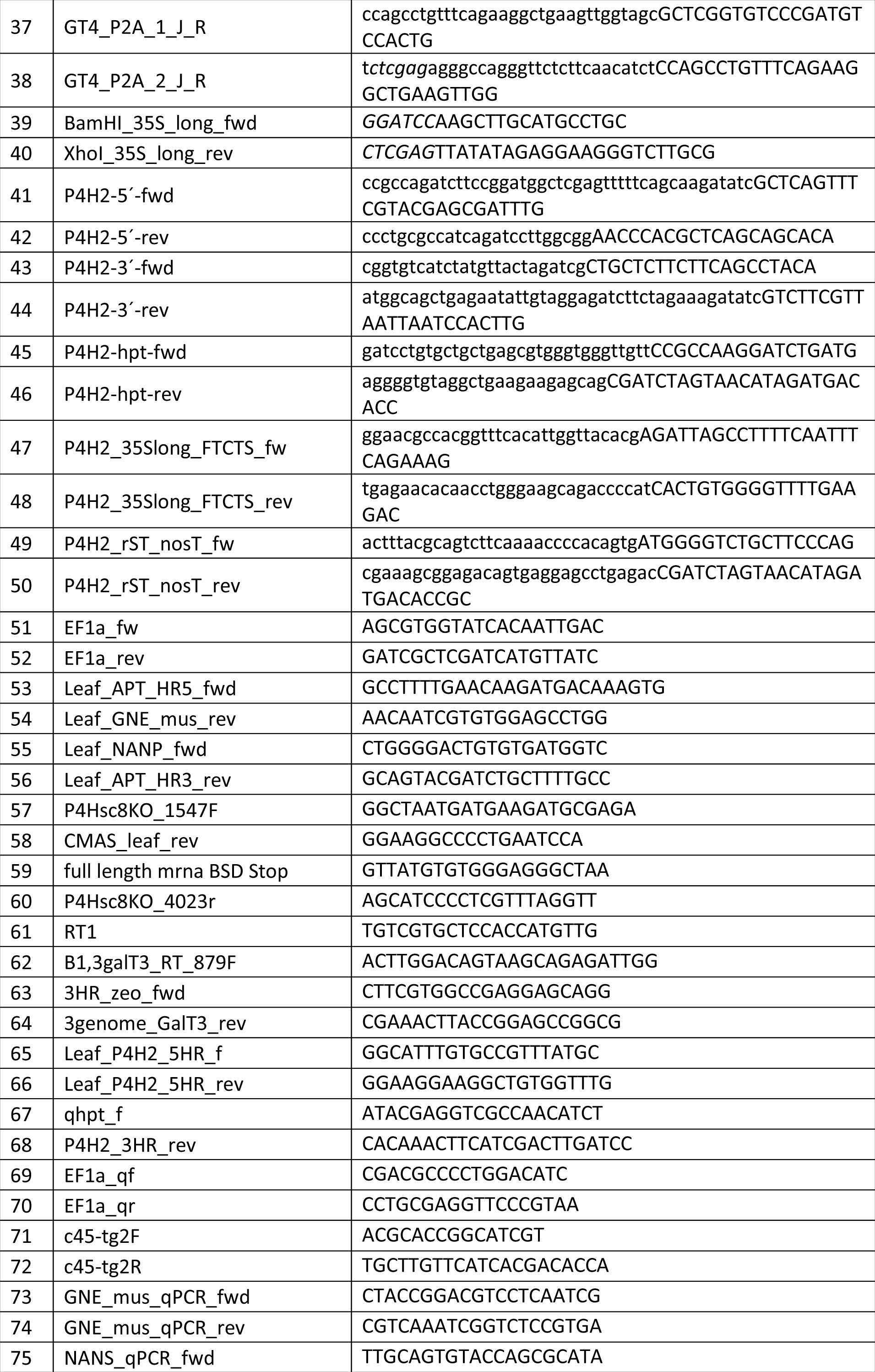
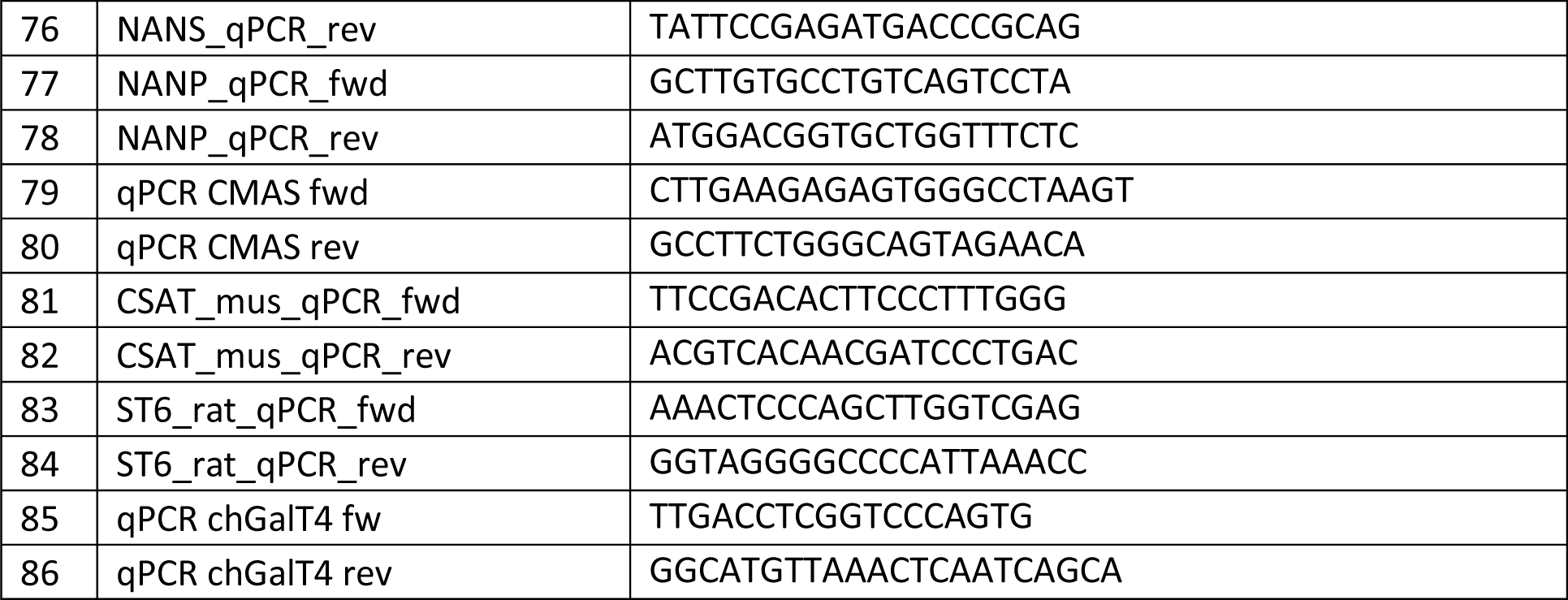
Primers used. Introduced restriction sites are written in italic. Lower-case letters indicate sequences of primer overhangs. Bolt letters indicate sequence mismatches for site-directed mutagenesis.

**Supplementary Table S2.**
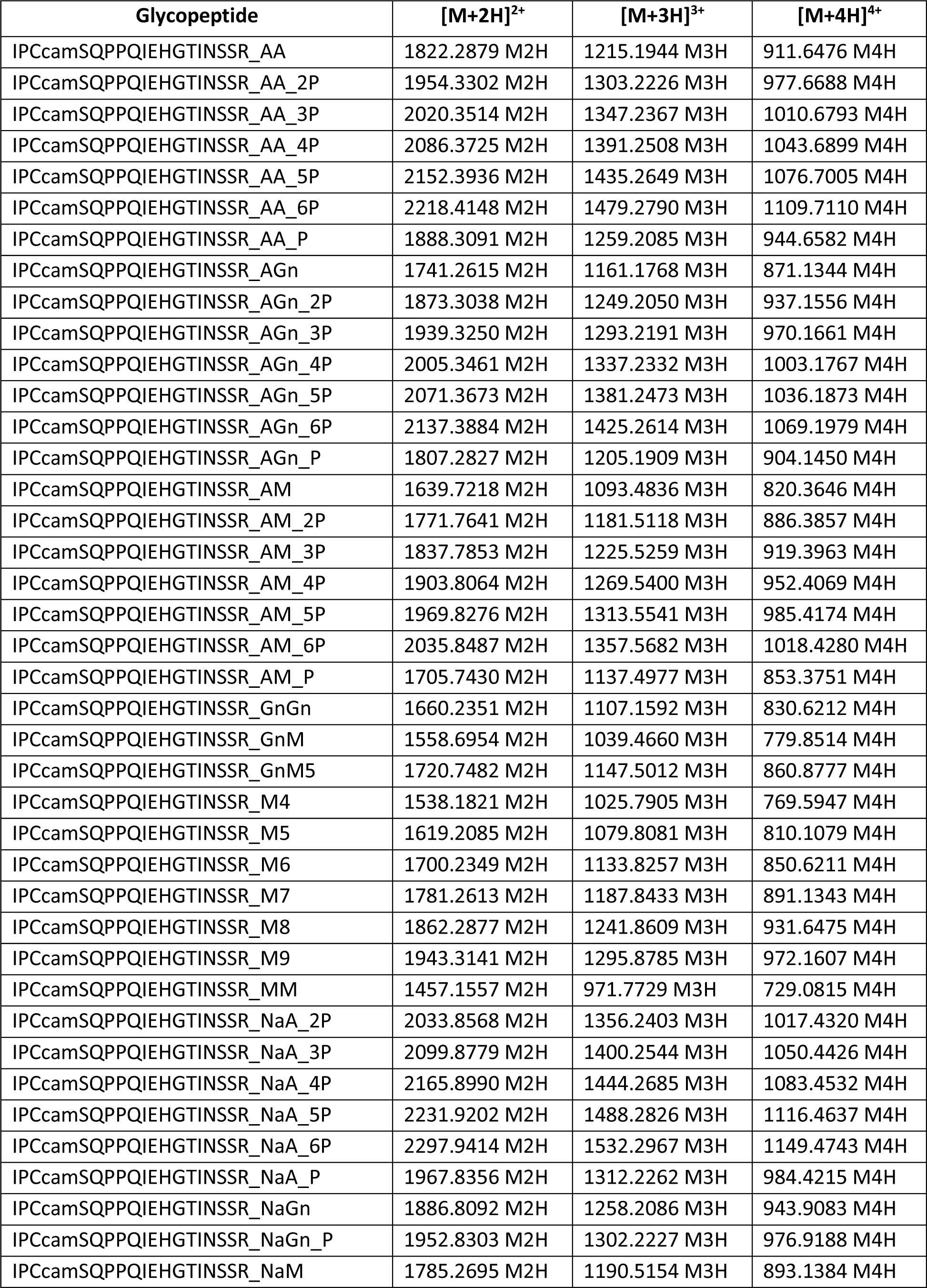
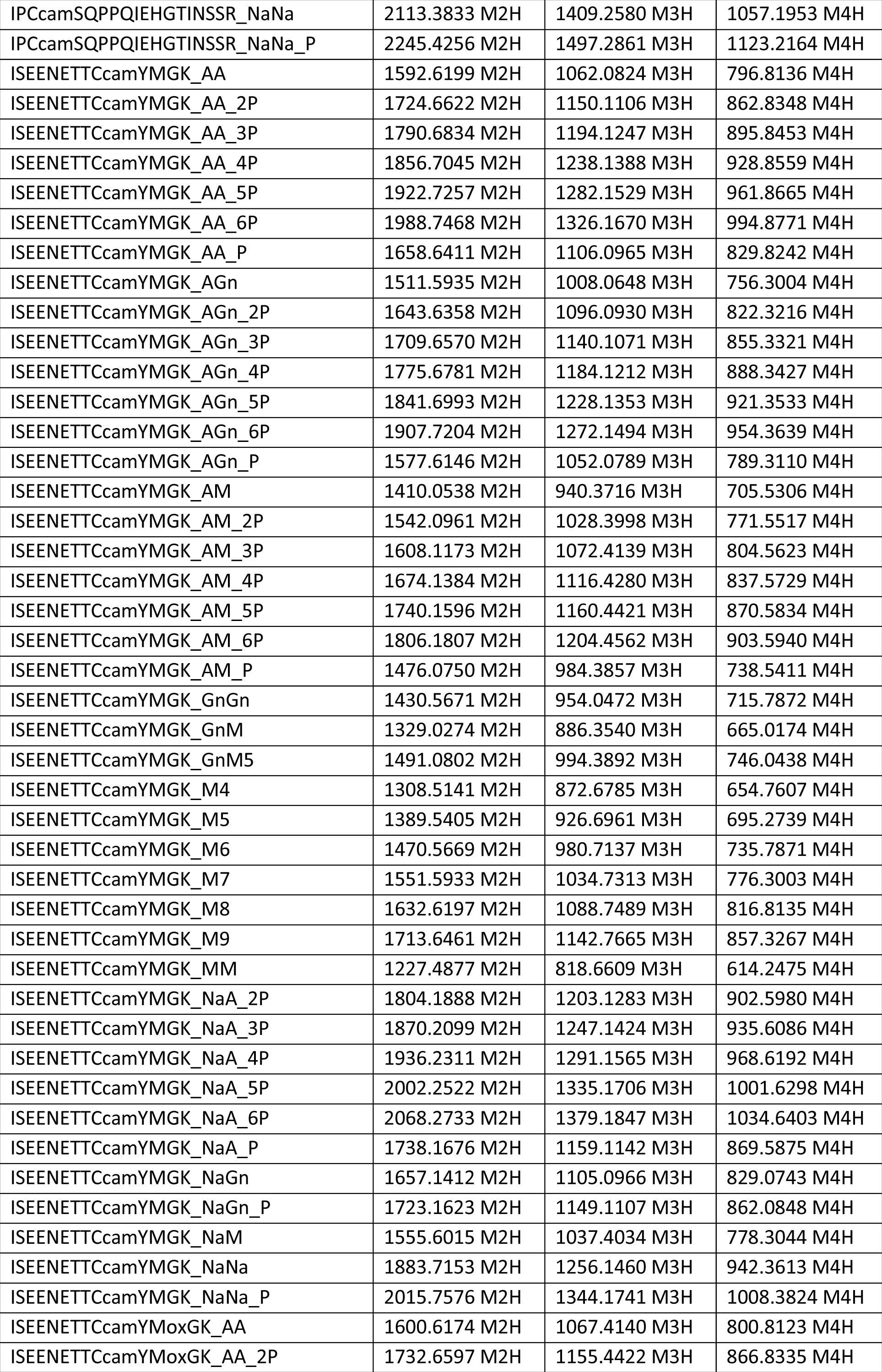
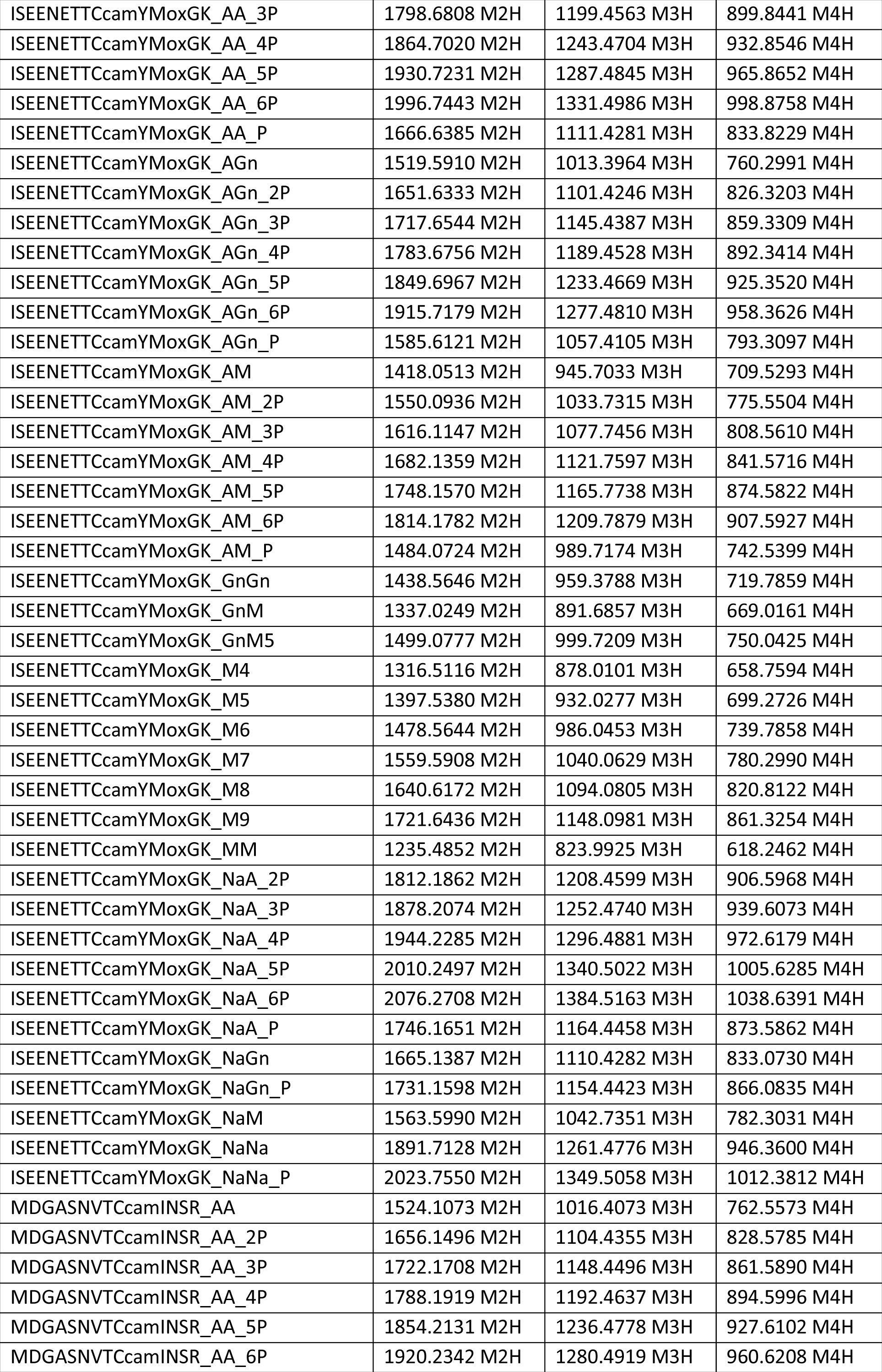
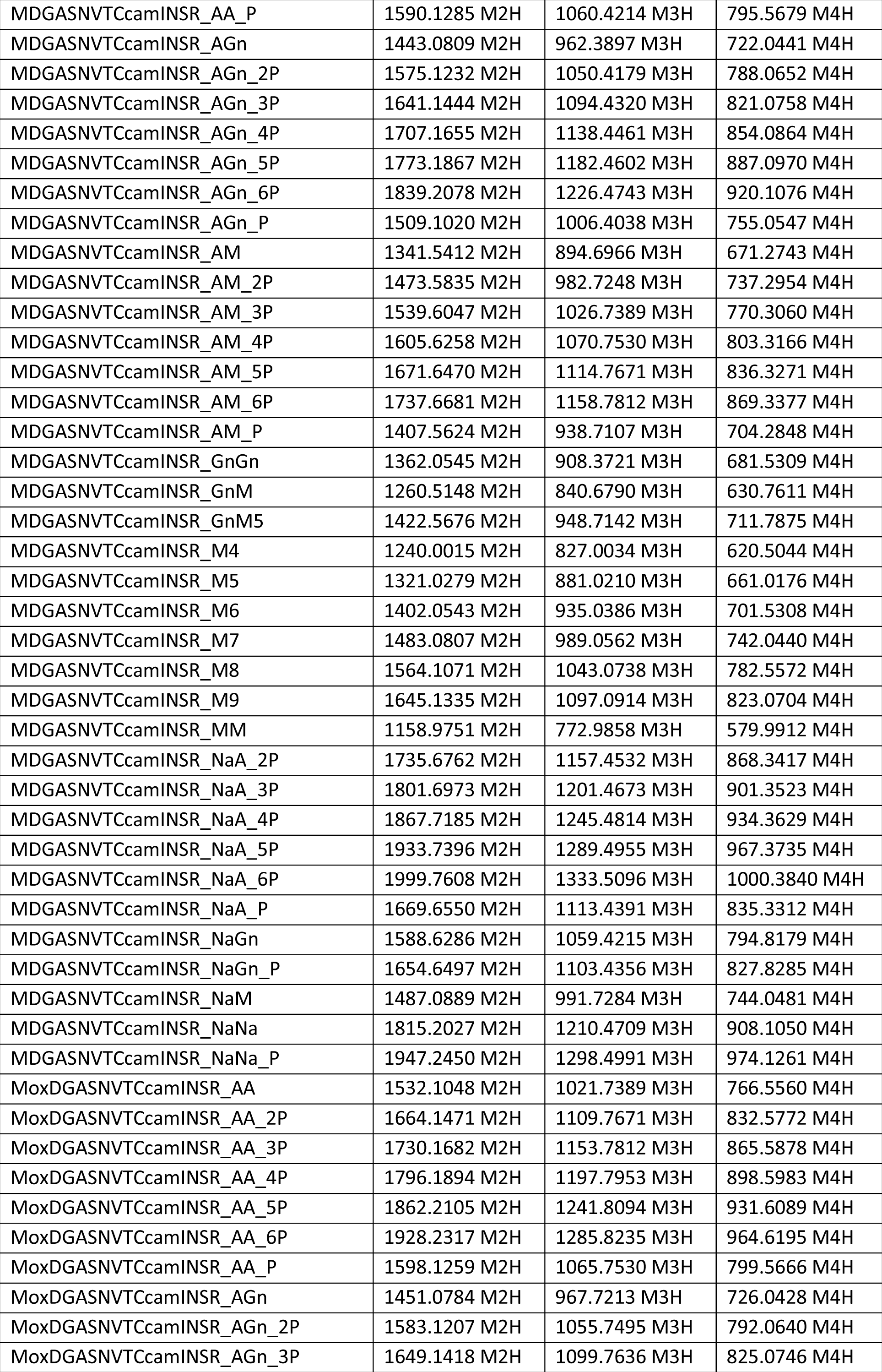
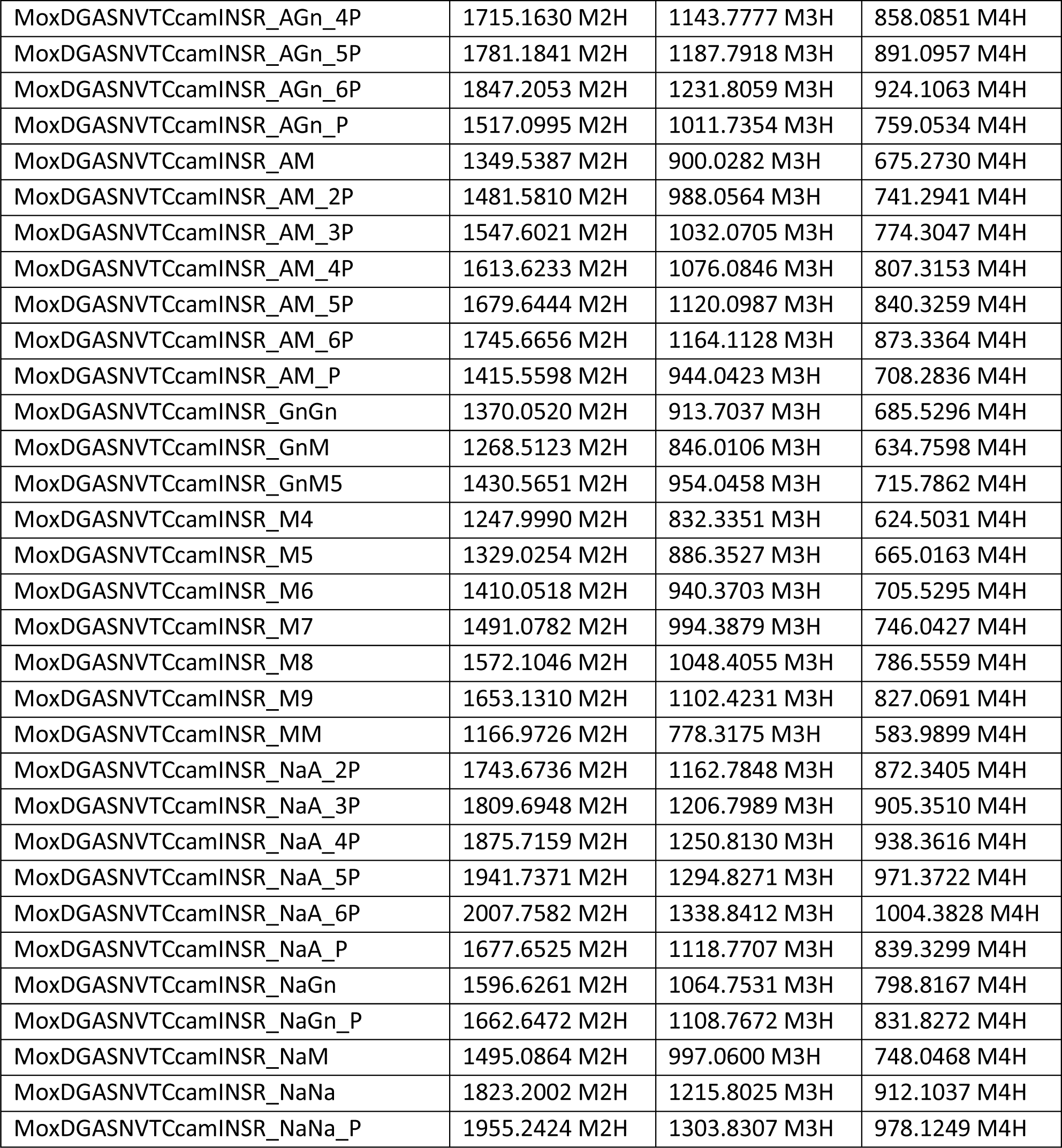
Monoisotopic masses of glycopeptides used for mass spectrometric analysis. All combinations of the searched peptides with possible glycan structures are given in three calculated charge stages ([M+nH]^n+^). cam: carbamidomethylation (C, + 57.021464 Da, fixed); ox: oxidation (M, +15.994915 Da, optional); M: mannose, Gn: *N*-acetylglucosamine, A: galactose, Na: sialic acid, P: pentose.

**Supplementary Figure S1.**
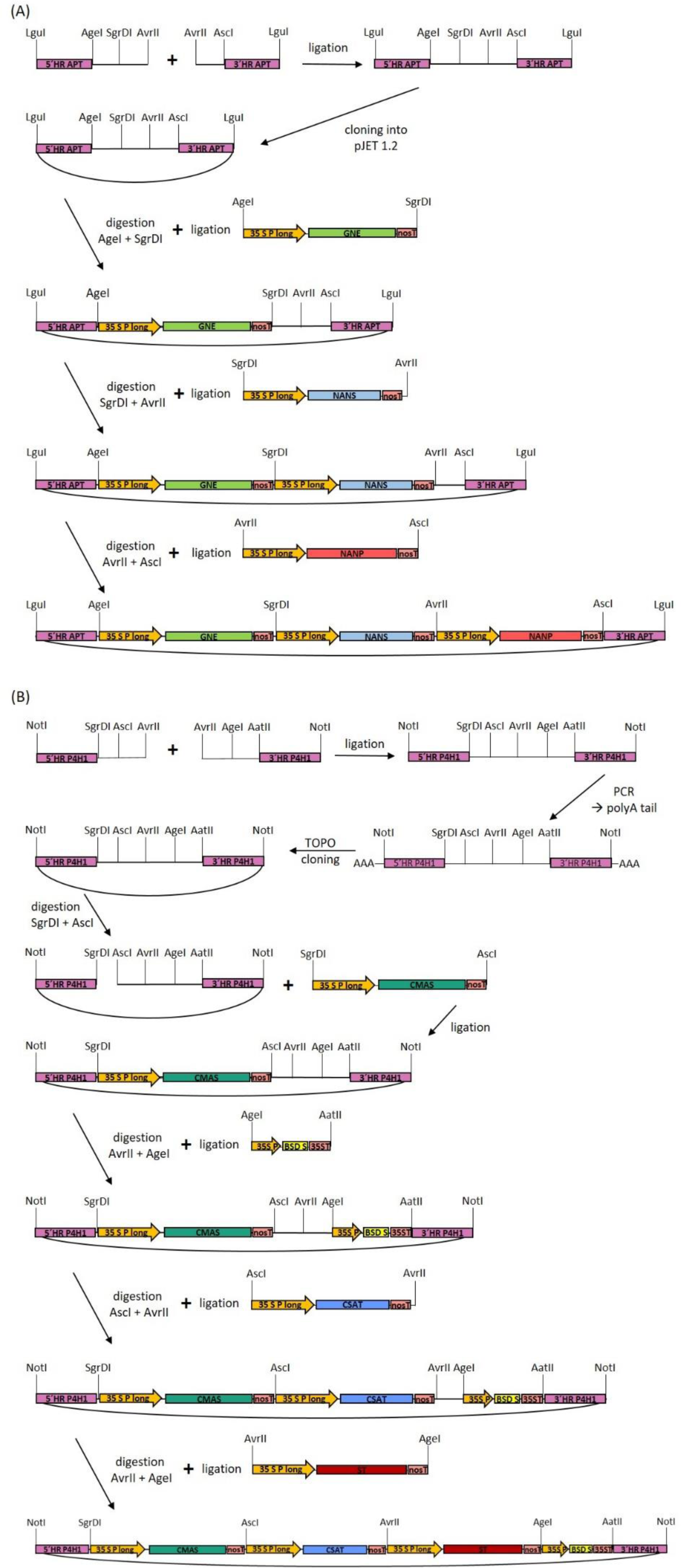
Schematic overview of the cloning procedure for the GNN and CCSB multi-gene constructs. **(A)** Successive cloning procedure of the GNN-construct starting with the generation of the pJET1.2-based assembly vector with a designed multiple cloning site and homologous flanks for integration into the adenine phosphoribosyltransferase (APT, Pp3c8_16590V3.1) locus. This vector was subsequently linearized with two restriction enzymes cutting within the designed MCS and ligated with the respective transgene expression cassette digested with the same enzymes from the expression vector. This was performed subsequently for the GNE-, NANS, and NANP-expression cassettes, resulting in the assembled GNN-construct. **(B)** Successive cloning procedure of the CCSB-construct starting with the generation of the pTOPO-based assembly vector with a designed multiple cloning site and homologous flanks for integration into the prolyl-4-hydroxylase 1 (P4H1, Pp3c8_7140V3.1) locus. This vector was subsequently linearized with two restriction enzymes cutting within the designed MCS and ligated with the respective transgene expression cassette digested with the same enzymes from the expression vector. This was performed subsequently for the CMAS, CSAT, ST and BSD-expression cassettes, resulting in the assembled CCSB-construct. The restriction enzymes used are indicated above the constructs. 5’HR: 5’-homologous region, 3’HR: 3’-homologous region, 35SP_long: long CaMV 35S promoter, 35SP: CaMV 35S promoter, nosT: nos terminator, 35ST: CaMV 35S terminator.

**Supplementary Figure S2.**
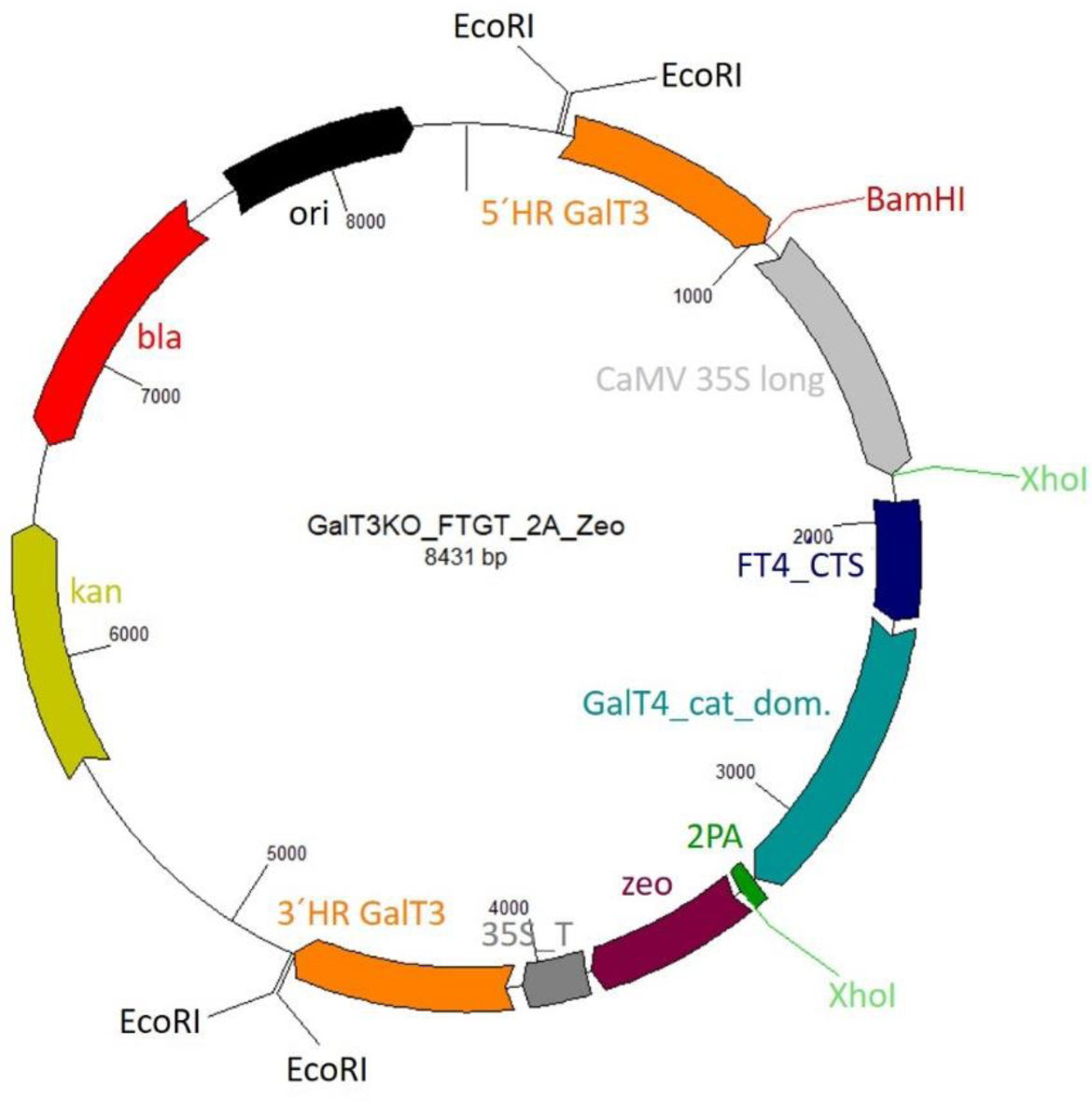
Schematic illustration of the plasmid containing the FTGT expression construct for targeted integration into the GalT3 locus. The homologous 5’- and 3’-flanks (5’HR GalT3, 3’HR GalT3) for targeted integration into the β1,3-galactosyltransferase 1 (GalT3) locus are displayed in orange. CaMV 35S long: long CaMV 35S promoter, FT4_CTS: sequence encoding the N-terminus of moss α1,4-fucosyltransferase including the CTS domain; GalT4_cat_dom.: sequence encoding the catalytic domain of the human GalT4, P2A: 2A peptide sequence from the porcine teschovirus-1, zeo: zeocine resistance coding for bleomycin resistance protein, 35S_T:CaMV 35S terminator, kan: kanamycin selection cassette, bla: ampicillin resistance cassette coding for beta-lactamase.

**Supplementary Figure S3.**
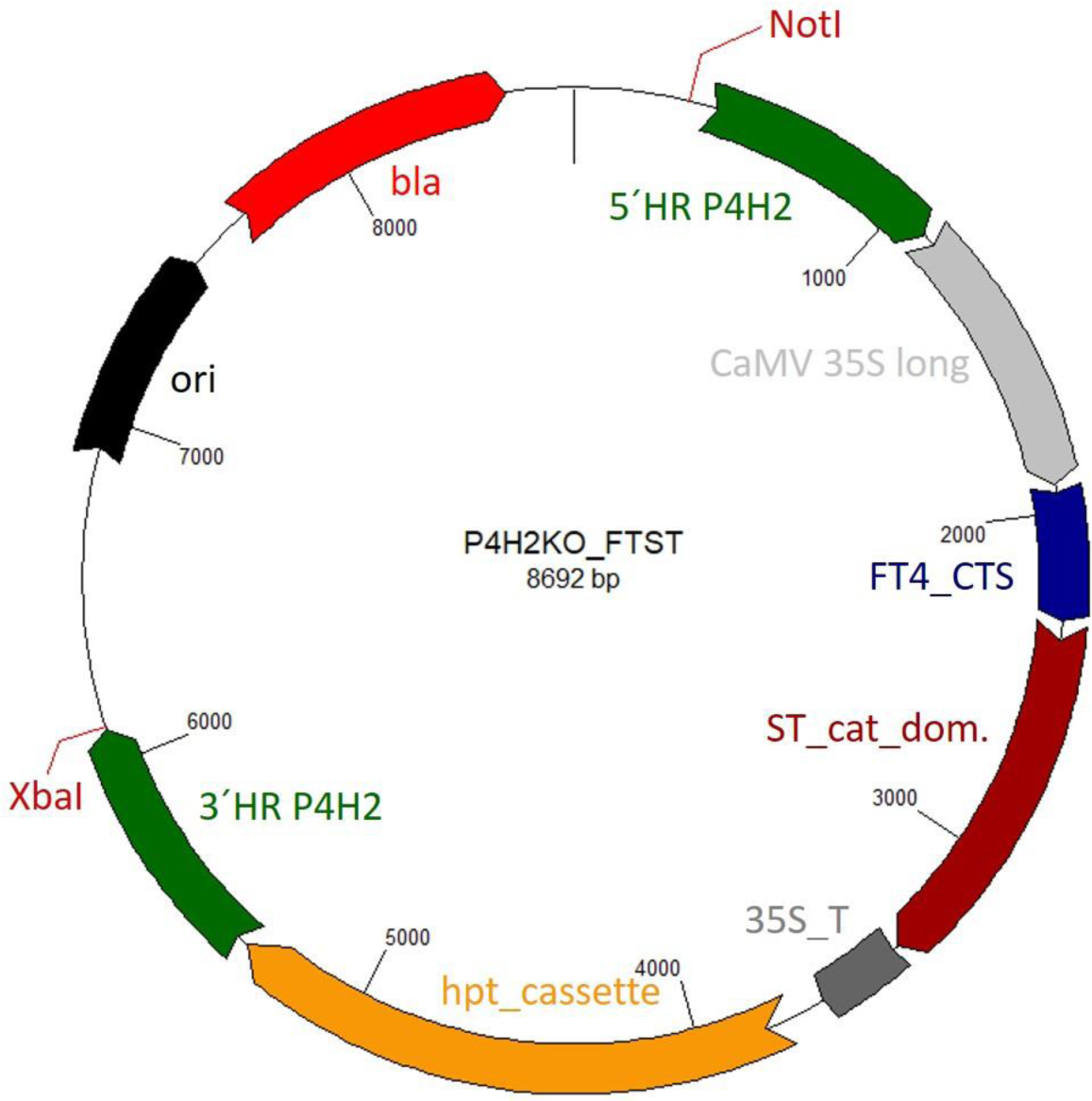
Schematic illustration of the plasmid containing the FTST expression construct for targeted integration into the P4H2 locus. The homologous 5’- and 3’-flanks (5’HR_P4H2, 3’HR_P4H2) for targeted integration into the prolyl-4-hydroxylase 2 (P4H2) locus are displayed in green. CaMV 35S long: long CaMV 35S promoter, FT4_CTS: sequence encoding the N-terminus of moss α1,4-fucosyltransferase including the CTS domain; ST_cat_dom.: sequence encoding the catalytic domain of the rat α2,6-sialyltransferase, 35S_T: CaMV 35S terminator, hpt_cassette: hygromycin B phosphotransferase encoding resistance gene driven by the nos promotor and nos terminator, bla: ampicillin resistance cassette coding for beta-lactamase.

**Supplementary Figure S4.**
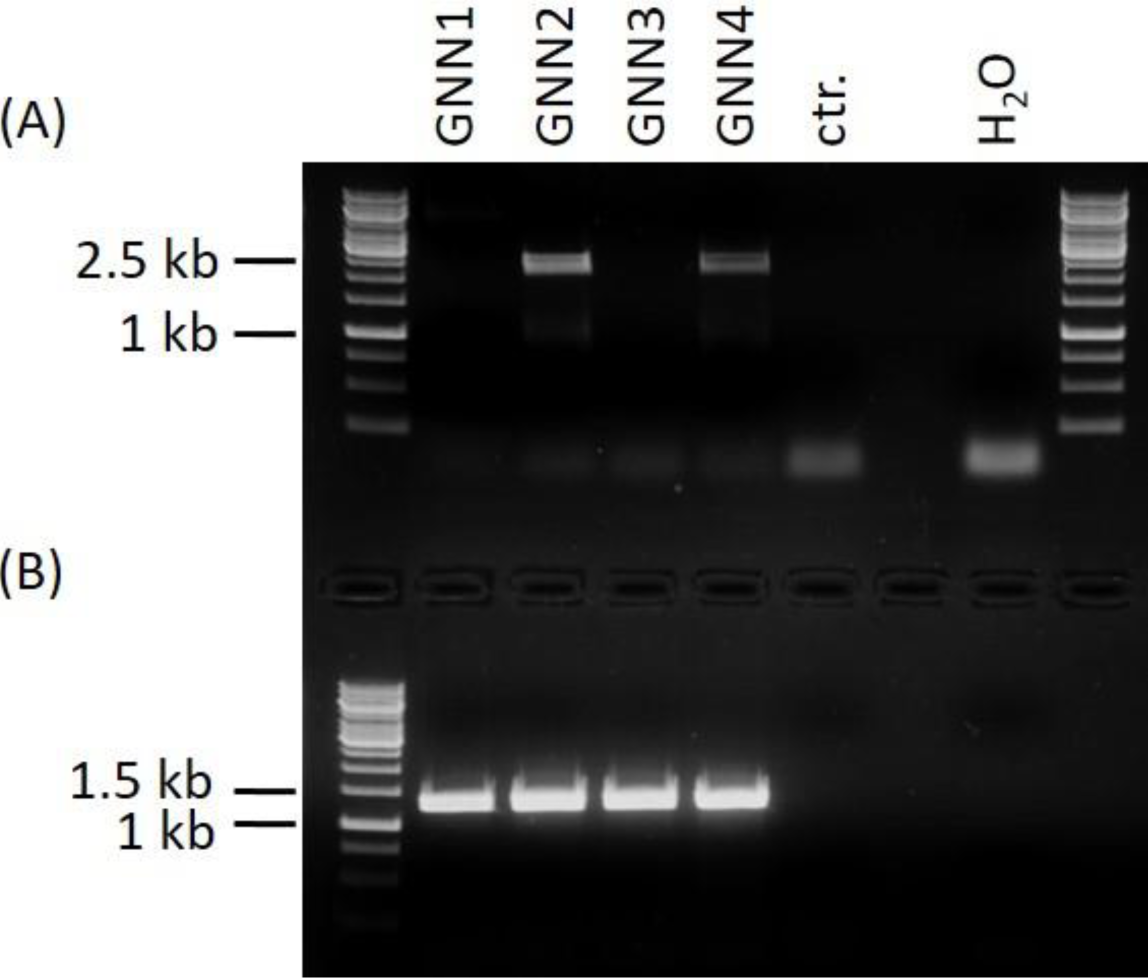
PCR-based analysis of 5’- (A) and 3’-integration (B) of the GNN-construct within the APT locus. PCR reactions were performed using primer pairs 53 and 54 for 5’- and 55 and 56 for 3’-integration on genomic DNA of lines GNN1-4 as well as of parental line as negative control (ctr.). To demonstrate the absence of contamination a control without DNA (H_2_O) was included. Amplicons of 2244 bp for the 5’- (**(A)**,GNN2 and GNN4) and 1311 bp for the 3’-integration **(B)** confirmed the targeted integration.

**Supplementary Figure S5.**
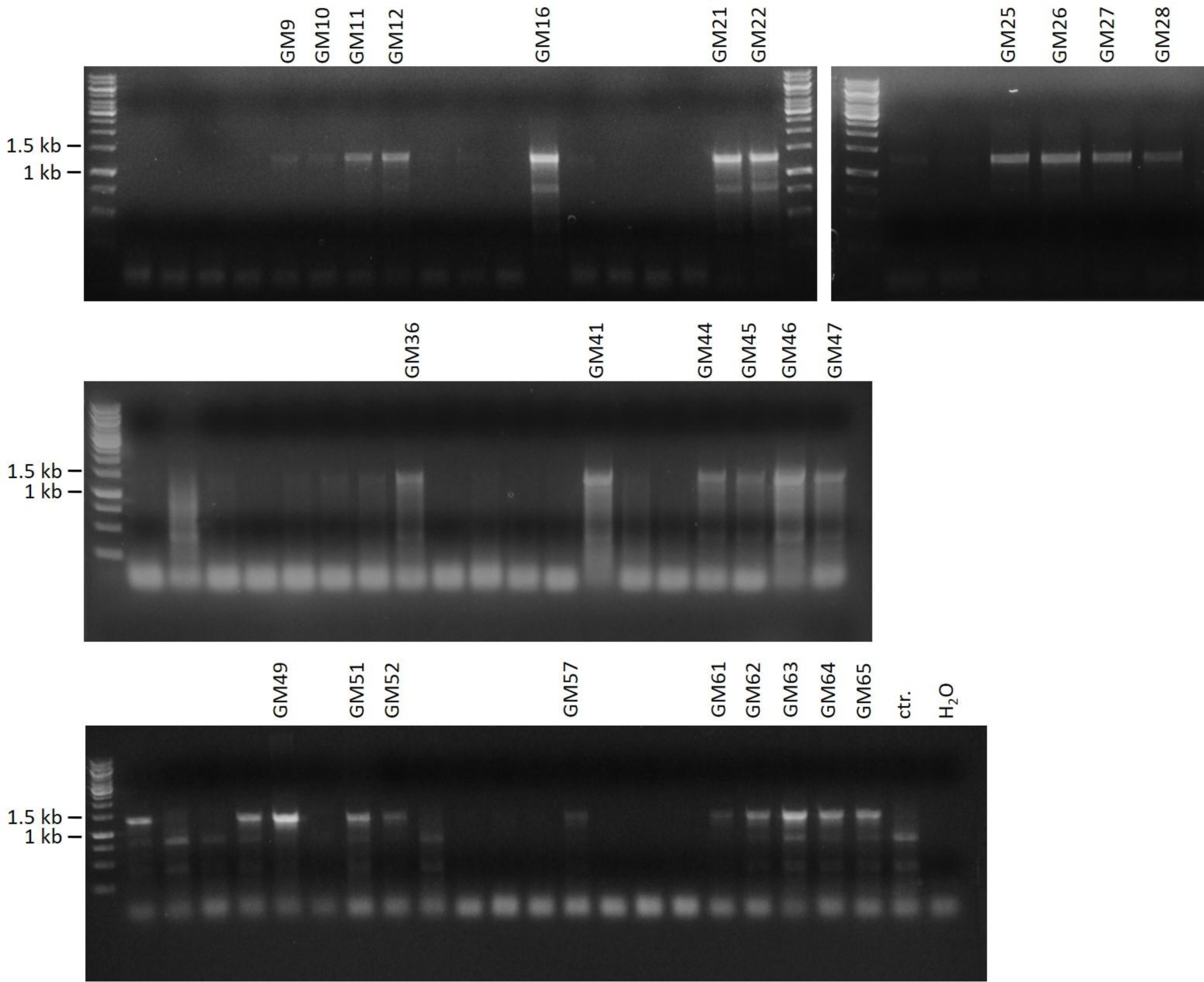
PCR-based analysis of 3’-integration of the GM-construct within the APT locus. PCR reactions were performed using the primer pairs 55 and 56 for 3’-integration on genomic DNA of lines GM1-65 as well as of a negative control (ctr.). To demonstrate the absence of contamination a control without DNA (H_2_O) was included. Amplicons of 1311 bp confirmed targeted 3’ construct integration.

**Supplementary Figure S6.**
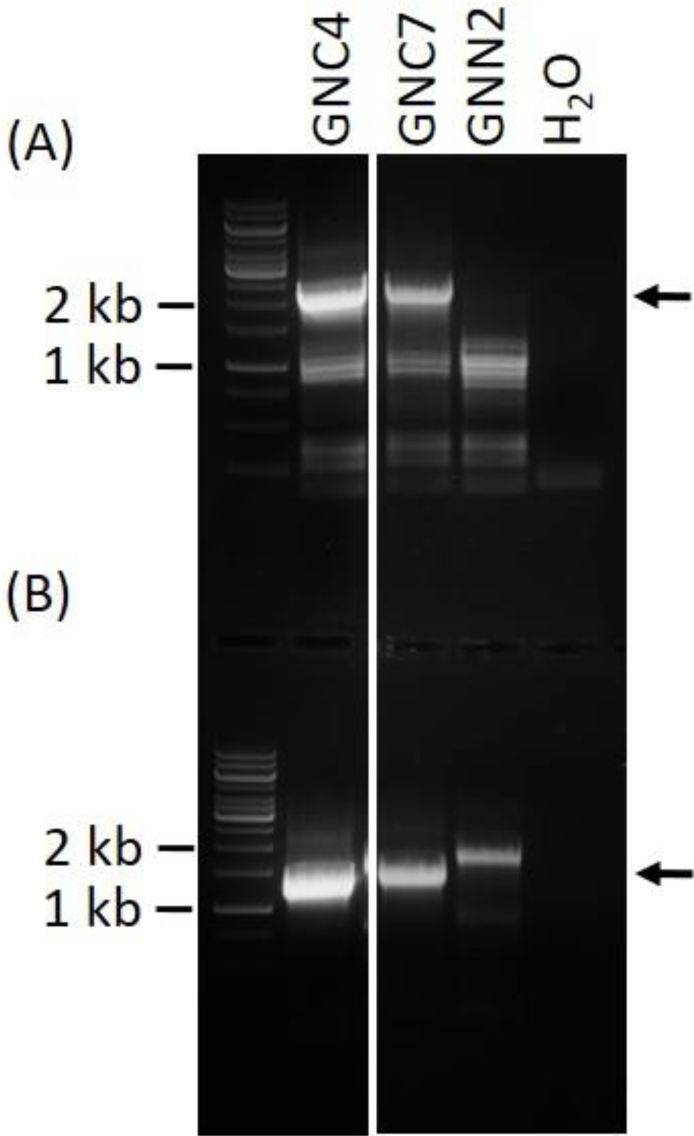
PCR-based analysis of 5’- and 3’-integration of the CCSB-construct in the P4H1 locus. PCR reactions were performed using primer pairs 57 and 58 for 5’- and 59 and 60 for 3’-integration on genomic DNA of lines GNC4 and GNC7 as well as of the GNN2-parental line as negative control. To demonstrate the absence of contamination a control without DNA (H_2_O) was included. Amplicons of 2222 bp for the 5’- **(A)** or 1406 bp for the 3’-integration **(B)** confirmed the targeted integration in the lines GNC4 and GNC7 (indicated by the black arrows).

**Supplementary Figure S7.**
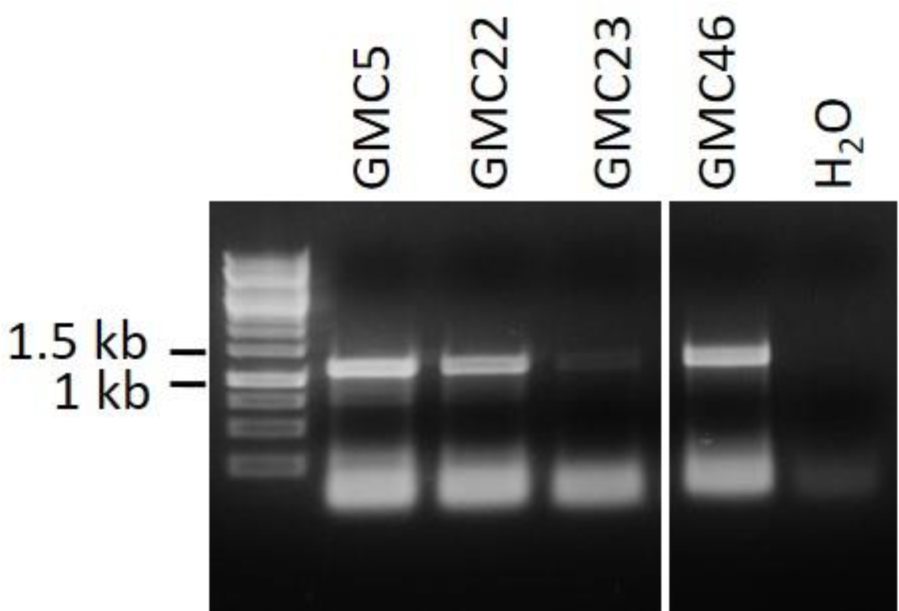
PCR-based analysis of 3’-integration of the CCSB-construct in the P4H1 locus. PCR reactions were performed using primer pair 59 and 60 for 3’-integration on genomic DNA of lines GMC5, 22, 23, and 46, resulting from CCSB transfections of line GM28. To demonstrate the absence of contamination a control without DNA (H_2_O) was included. Amplicons of 1406 bp confirmed targeted 3’ construct integration in all analyzed lines.

**Supplementary Figure S8.**
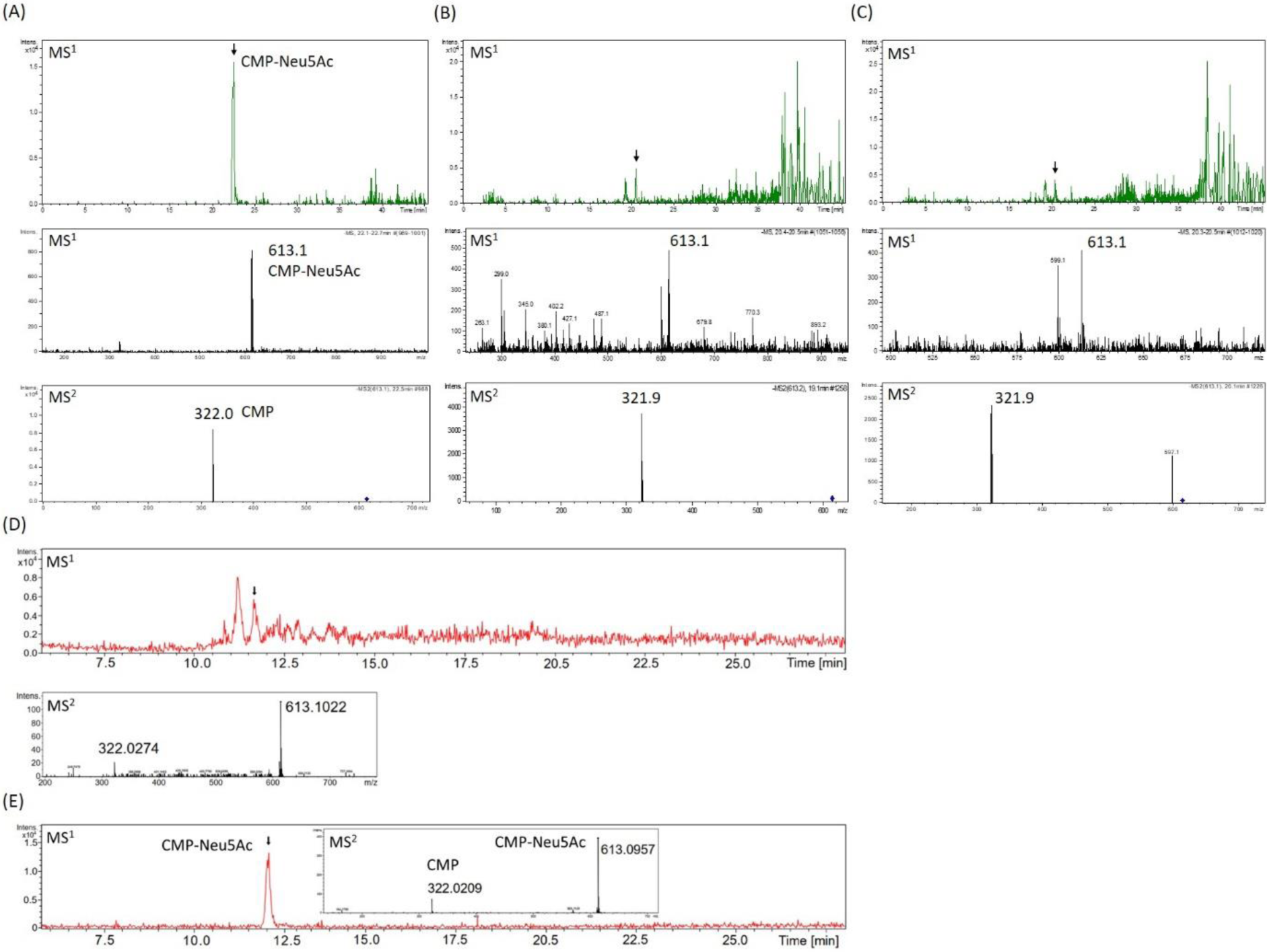
Mass-spectrometric detection of CMP-Neu5Ac in moss extracts **(B-D)** compared to measurements of a CMP-Neu5Ac standard **(A+E)**. Extracted ion chromatogram (m/z of 613.1 ± 0.1) of LC-MS measured extracts of the lines GMC23 **(B)**, GMC46 **(C)**, GNC7 **(D)** and 70 pmol of a CMP-Neu5Ac standard (**A** (standard measurement for B + C) and **E** (standard measurement for D)), respectively (upper panels A-D and E). In all measurements peaks corresponding to the [M-H]^−^-ion of CMP-Neu5Ac of m/z = 613.1 could be detected on MS^1^ level (middle panels A-C, E and lower panel of D). Confirmation of corresponding MS^1^-peak identities (indicated by the black arrows) was performed on MS^2^ level via the identification of the [M-H]^-^ CMP fragment ion of m/z = 322.0 (lower panels A-D and E). CMP-Neu5Ac concentration of the moss extracts was determined via peak area integration in comparison to defined standard values and resulted in 14 nmol CMP-Neu5Ac/g DW in the lines GMC23 and 46 and 2.3 nmol CMP-Neu5Ac/g DW in GNC7.

**Supplementary Figure S9.**
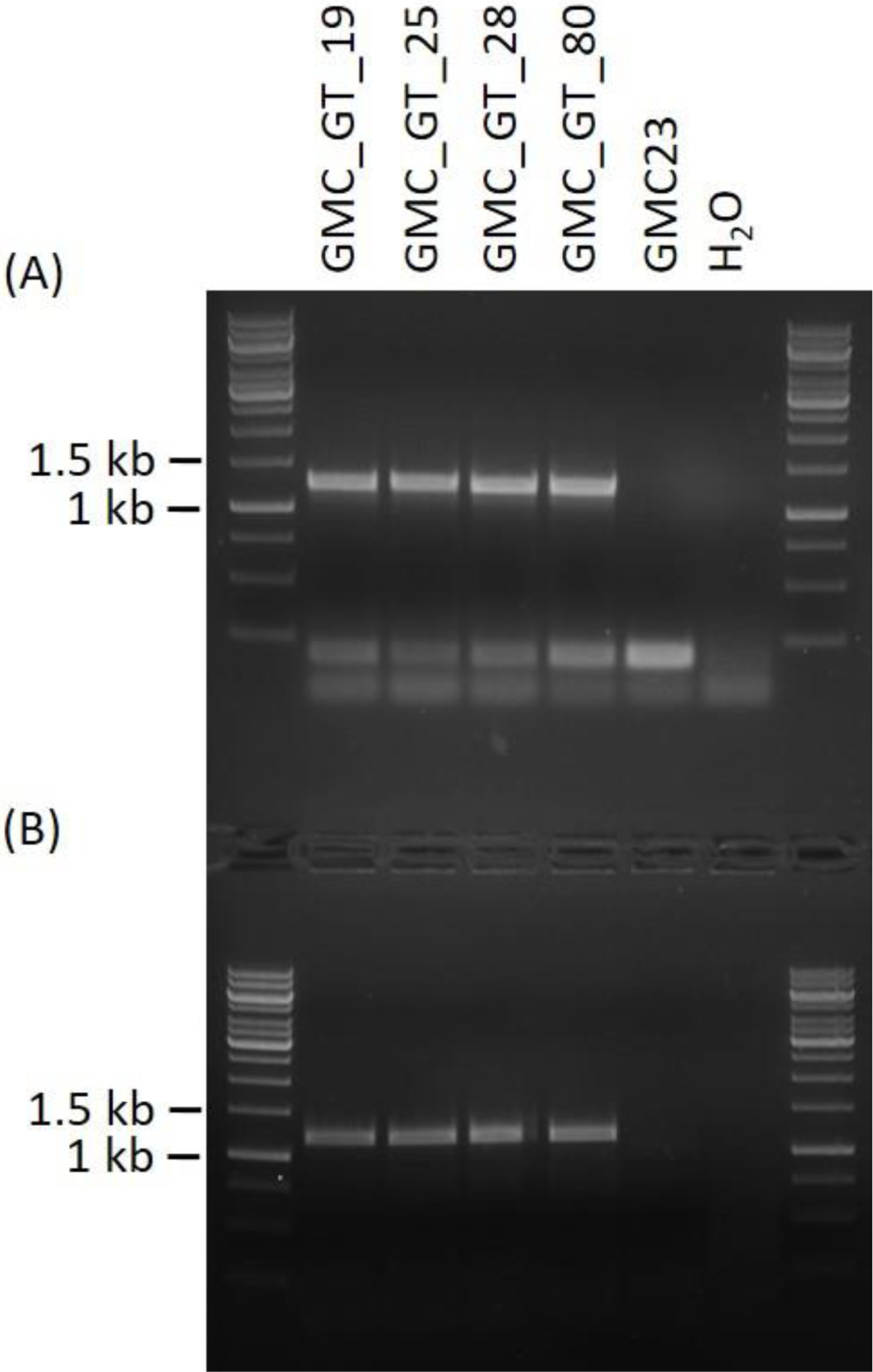
PCR-based analysis of 5’-and 3’-integration of the FTGT-construct in the GalT3 locus. PCR reactions were performed on genomic DNA of the lines GMC_GT_19, 25, 28 and 80 as well as the GMC23 parental line using the primer pairs 61 and 62 for 5’- as well as 63 and 64 for 3’-integration, respectively. To demonstrate the absence of contamination a control without DNA (H_2_O) was included. These analyses resulted in integration-confirming amplicons of 1253 for the 5’- **(A)** and 11171 bp for the 3’-integration **(B)** for all transgenic lines, whereas no integration-indicating signals were obtained in the parental line and the H_2_O control.

**Supplementary Figure S10.**
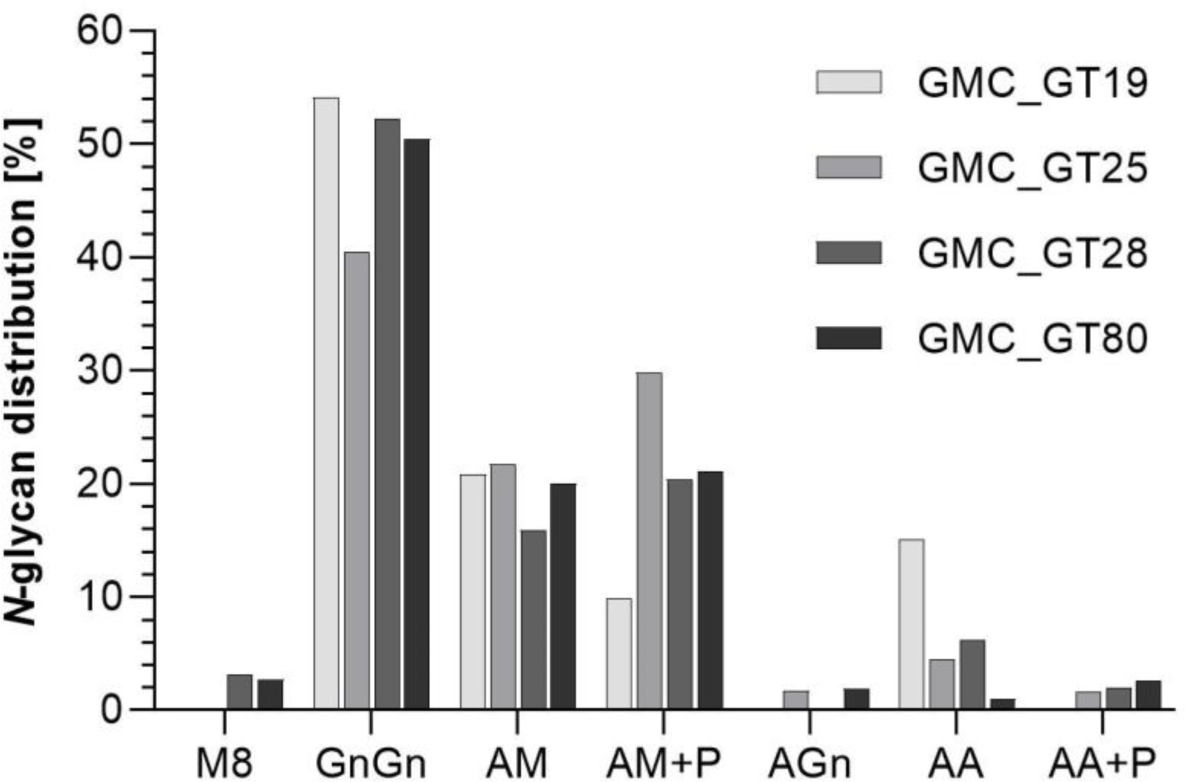
Mass-spectrometric analysis of *N*-glycosylation patterns in GMC_GT lines. MS-based relative quantification of identified tryptic glycopeptides from the reporter glycoprotein of complete sialyation lines. Relative quantification was based on peak area integration of extracted ion chromatograms on MS^1^ level, for which peak identities were confirmed on MS^2^ level. For quantification, areas of all confirmed peaks per measurement were summed up and the relative percentages are given for each identified glycan structure. M: mannose, Gn: *N*-acetylglucosamine, A: galactose, P: represents the presence of one or two pentoses attached to the *N*-glycan.

**Supplementary Figure S11.**
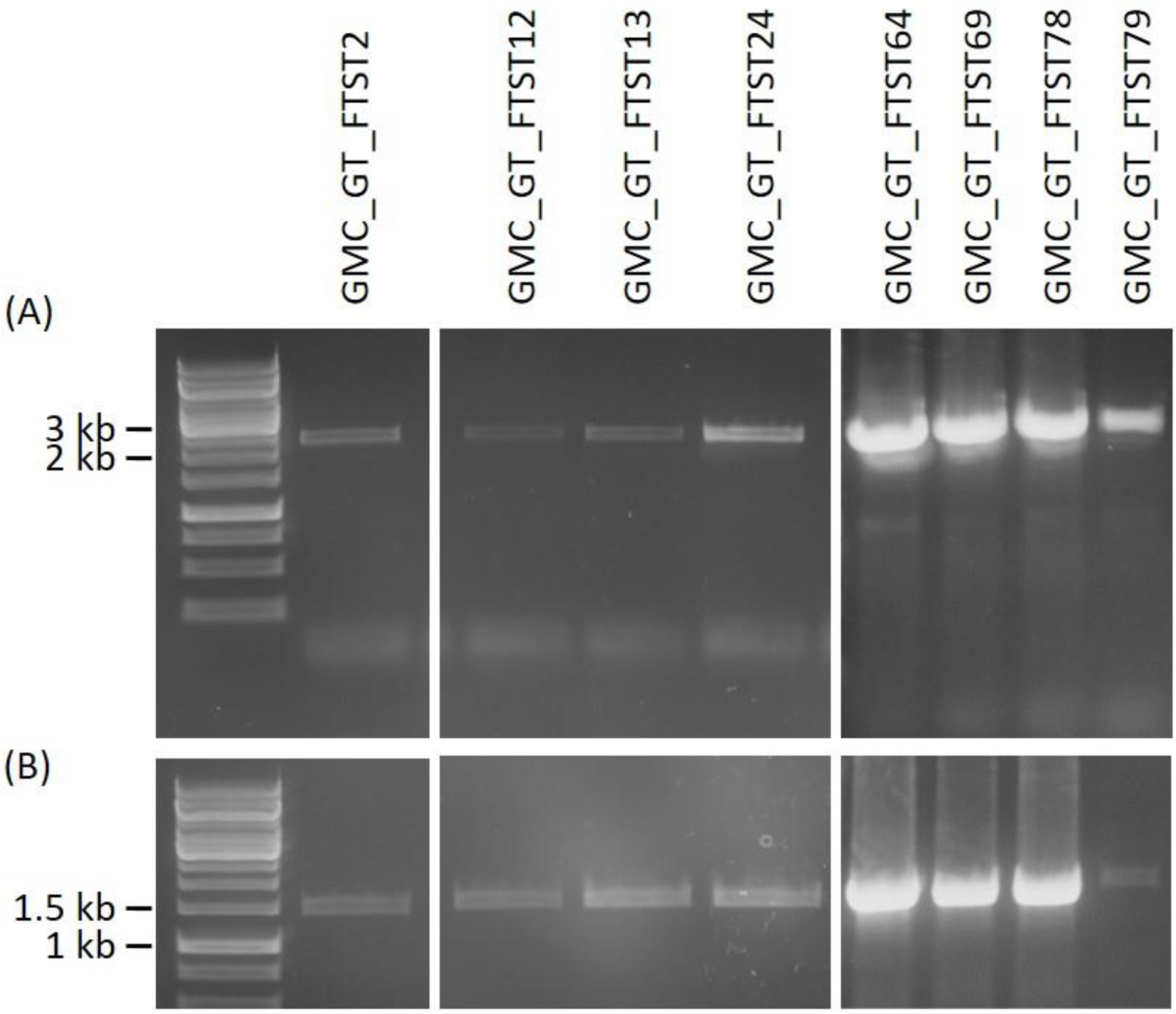
PCR signals confirming targeted 5’-and 3’-integration of the FTST-construct within the P4H2 locus. PCR reactions were performed on genomic DNA using the primer pairs 65 and 66 for 5’- as well as 67 and 68 for 3’-integration, respectively. These analyses resulted for all analyzed lines in integration-confirming amplicons of 2169 for the 5’- **(A)** and 1464 bp for the 3’-integration **(B)**, respectively.

**Supplementary Figure S12.**
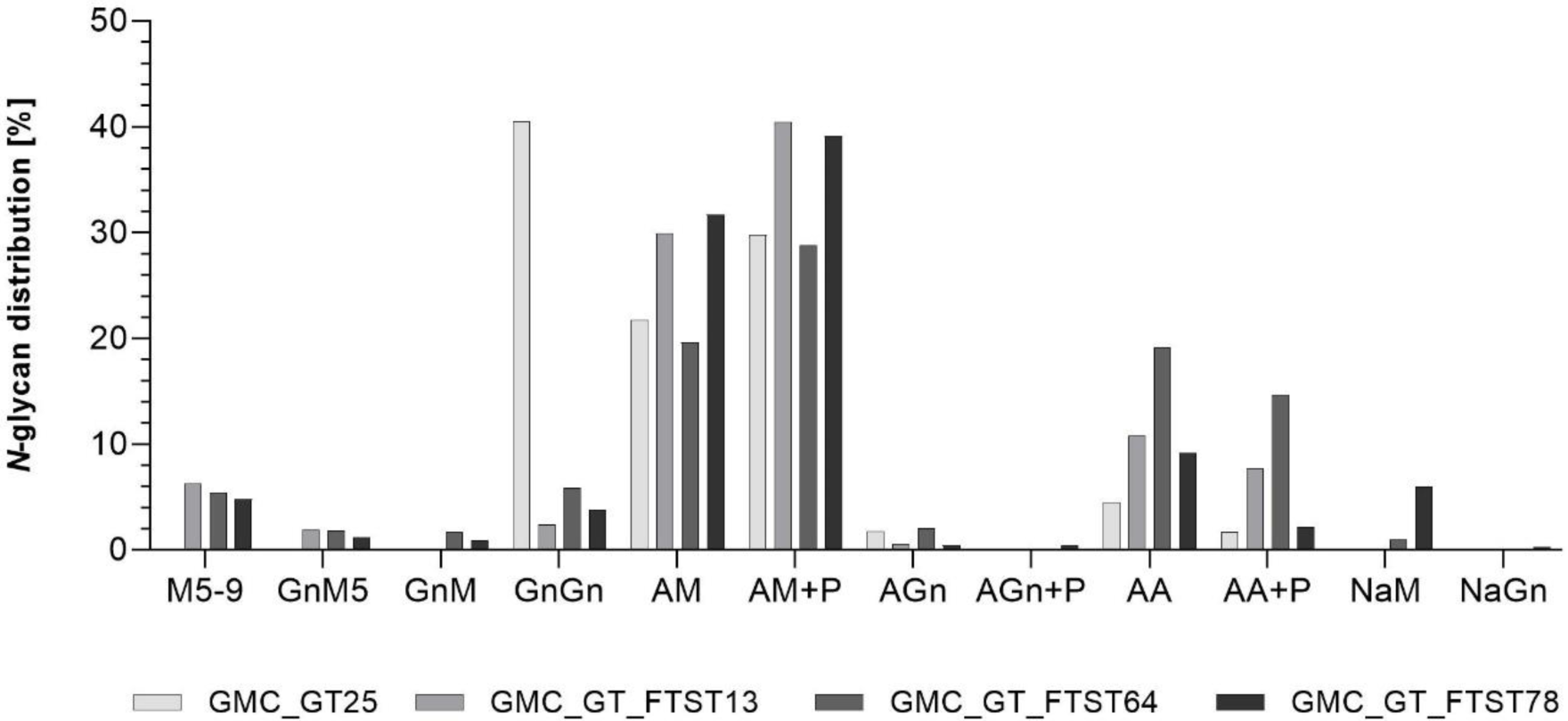
Mass-spectrometric analysis of glycosylation patterns in FTST-expressing lines compared to the GMC_GT25 parental line. MS-based relative quantification of glycopeptides identified on the tryptically digested reporter glycoprotein. Relative quantification was based on peak area integration of extracted ion chromatograms on MS^1^ level, for which peak identities were confirmed on MS^2^ level. For quantification, areas of all confirmed peaks per measurement were summed up and the relative percentages are given for each identified glycan structure. M: mannose, Gn: *N-*acetylglucosamine, A: galactose, Na: sialic acid, P: indicates the presence of one or two pentoses on the corresponding *N-*glycan structure.

**Supplementary Figure S13.**
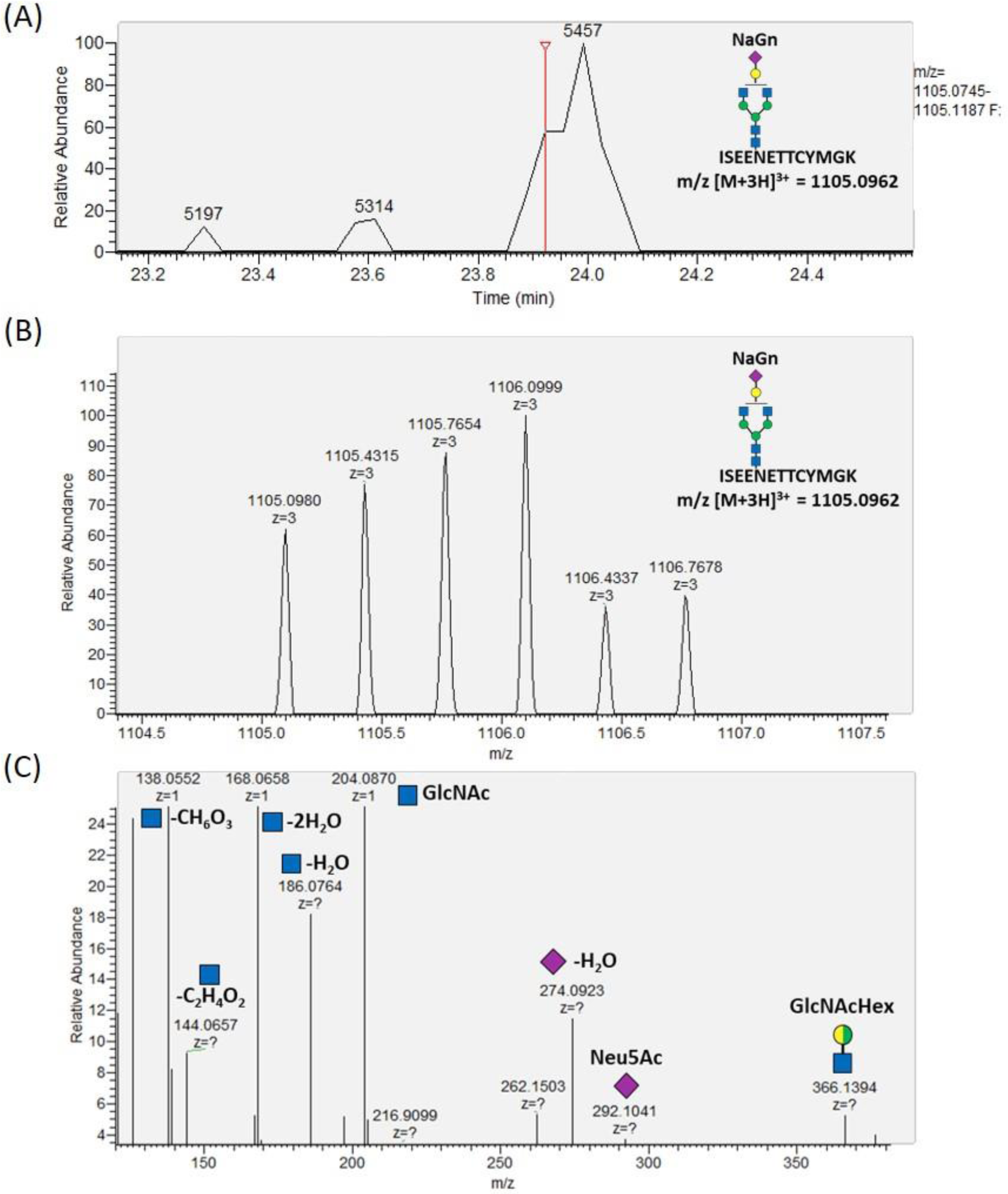
Mass-spectrometric verification of the identified NaGn-glycosylation of the ISEENETTCYMGK glycopeptide on MS^1^ **(A+B)** and MS^2^ level **(C). (A)** Extracted ion chromatogram of the expected [M+3H]^3+^ m/z-ratio of 1105.0962 of the tryptic NaGn-ISEENETTCYMGK glycopeptide on MS^1^ level. **(B)** Verification of the charge state (z=3) of the detected glycopeptide via investigation from the isotope pattern. **(C)** MS^2^-based verification of *N*-glycan sialylation via the detection of *N*-acetylglucosamine (GlcNAc) and sialic acid (Neu5Ac) reporter ions with the following m/z-values: [GlcNAc]^+^ = 204.087, [GlcNAc - H_2_O]^+^ = 186.076, [GlcNAc - 2H_2_O]^+^ = 168.066, [GlcNAc - C_2_H_4_O_2_]^+^ = 144.065, [GlcNAc - CH_6_O_3_]^+^ = 138.055, [GlcNAc - C_2_H_6_O_3_]^+^ = 126.055), [Neu5Ac]^+^ = 292.103, [Neu5Ac - H_2_O]^+^ = 274.092 and the detection of the glycan fragment ion [GlcNAcHex]^+^ = 366.139. Blue square: *N*-acetylglucosamine (Gn, GlcNAc), green circle: mannose (M), yellow circle: galactose (A), purple rhombus: sialic acid (Na, Neu5Ac), Hex: hexose (yellow and green circle: stands for the presence of either mannose or galactose).

**Supplementary Figure S14.**
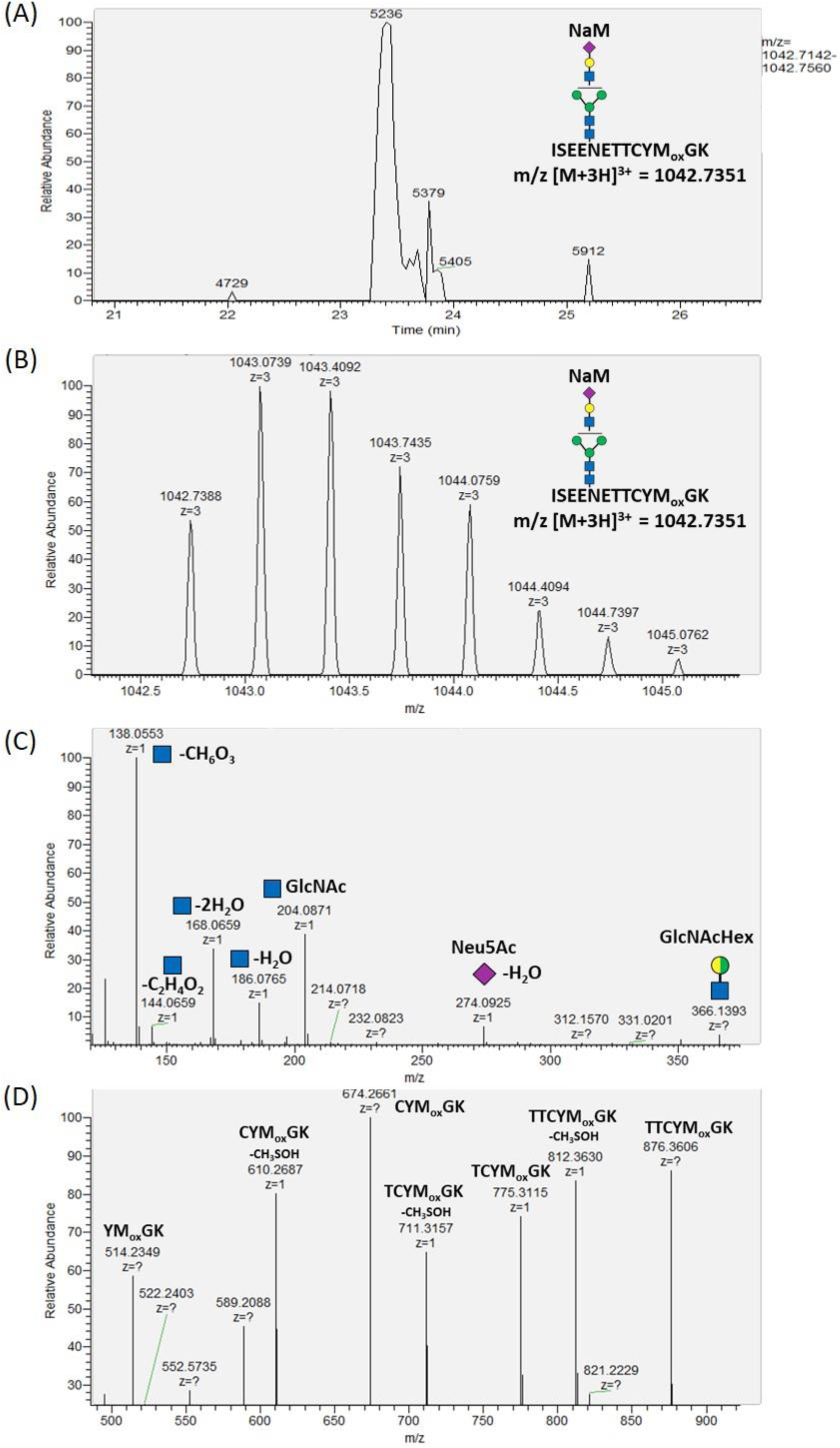
Mass spectrometric verification of the identified NaM-glycosylation of the ISEENETTCYM_ox_GK glycopeptide on MS^1^ **(A+B)** and MS^2^ level **(C+D). (A)** Extracted ion chromatogram of the expected [M+3H]^3+^ m/z-ratio of 1042.7351 of the tryptic NaM-ISEENETTCYM_ox_GK glycopeptide on MS^1^ level. **(B)** Verification of the charge state (z=3) of the detected glycopeptide via investigation from the isotope pattern. **(C)** MS^2^-based verification of *N*-glycan sialylation via the detection of *N*-acetylglucosamine (GlcNAc) and sialic acid (Neu5Ac) reporter ions with the following m/z-values: [GlcNAc]^+^ = 204.087, [GlcNAc - H_2_O]^+^ = 186.076, [GlcNAc - 2H_2_O]^+^ = 168.066, [GlcNAc - C_2_H_4_O_2_]^+^ = 144.065, [GlcNAc - CH_6_O_3_]^+^ = 138.055, [GlcNAc - C_2_H_6_O_3_]^+^ = 126.055), [Neu5Ac]^+^ = 292.103, [Neu5Ac - H_2_O]^+^ = 274.092 and the detection of the glycan fragment ion [GlcNAcHex]^+^ = 366.139. **(D)** Fragmentation of the peptide backbone. Blue square: *N*-acetylglucosamine (Gn, GlcNAc), green circle: mannose (M), yellow circle: galactose (A), purple rhombus: sialic acid (Na, Neu5Ac), Hex: hexose (yellow and green circle: stands for the presence of either mannose or galactose).

**Supplementary Figure S15.**
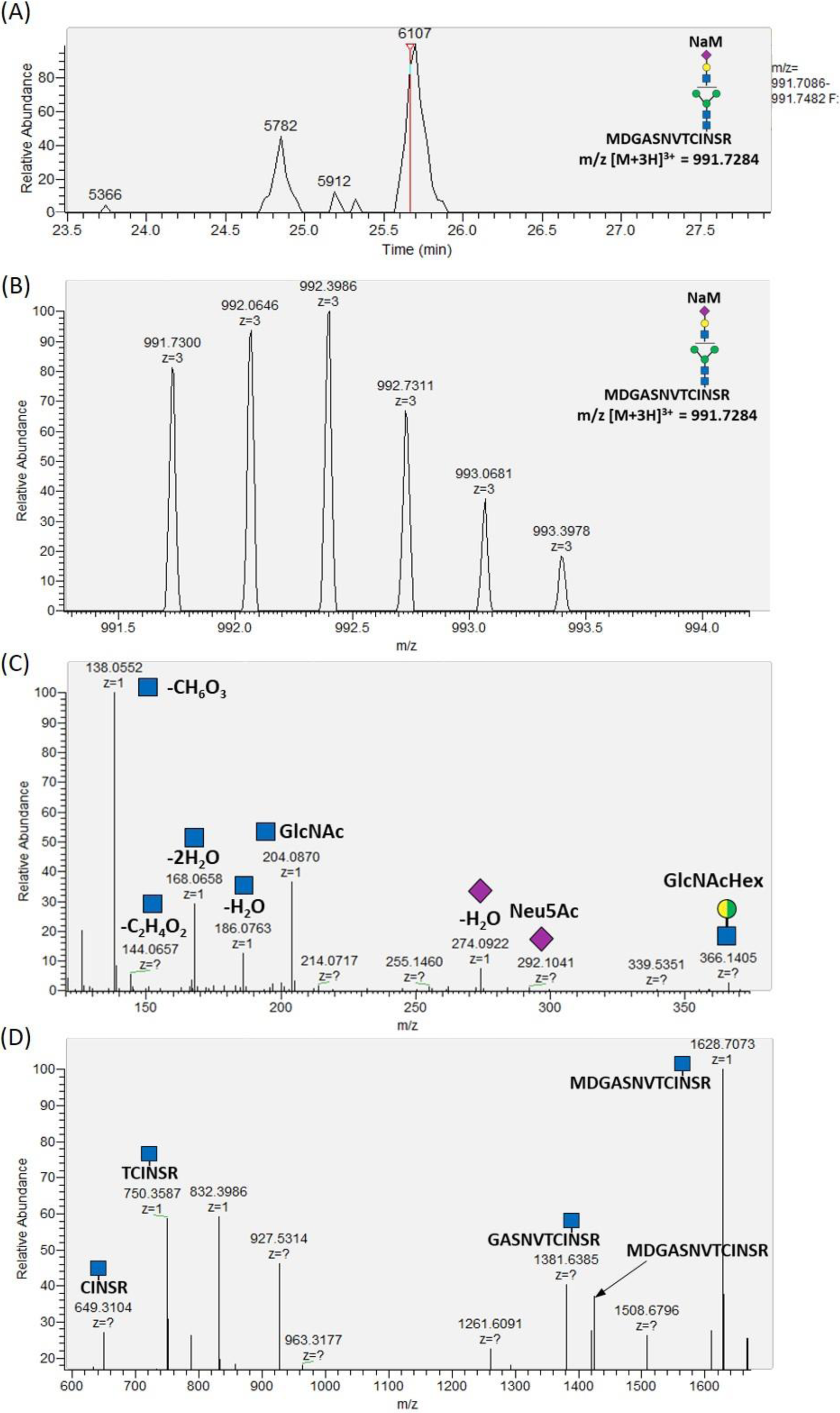
Mass-spectrometric verification of the identified NaM-glycosylation of the MDGASNVTCINSR glycopeptide on MS^1^ **(A+B)** and MS^2^ level **(C+D). (A)** Extracted ion chromatogram of the expected [M+3H]^3+^ m/z-ratio of 991.7284 of the tryptic NaM-MDGASNVTCINSR glycopeptide on MS^1^ level. **(B)** Verification of the charge state (z=3) of the detected glycopeptide via investigation from the isotope pattern. **(C)** MS^2^-based verification of *N*-glycan sialylation via the detection of *N-*acetylglucosamine (GlcNAc) and sialic acid (Neu5Ac) reporter ions with the following m/z-values: [GlcNAc]^+^ = 204.087, [GlcNAc - H_2_O]^+^ = 186.076, [GlcNAc - 2H_2_O]^+^ = 168.066, [GlcNAc - C_2_H_4_O_2_]^+^ = 144.065, [GlcNAc - CH_6_O_3_]^+^ = 138.055, [GlcNAc - C_2_H_6_O_3_]^+^ = 126.055), [Neu5Ac]^+^ = 292.103, [Neu5Ac - H_2_O]^+^ = 274.092 and the detection of the glycan fragment ion [GlcNAcHex]^+^ = 366.139. **(D)** Fragment ions of the glycopeptide. Blue square: *N-*acetylglucosamine (Gn, GlcNAc), green circle: mannose (M), yellow circle: galactose (A), purple rhombus: sialic acid (Na, Neu5Ac), Hex: hexose (yellow and green circle: stands for the presence of either mannose or galactose).

**Supplementary Figure S16.**
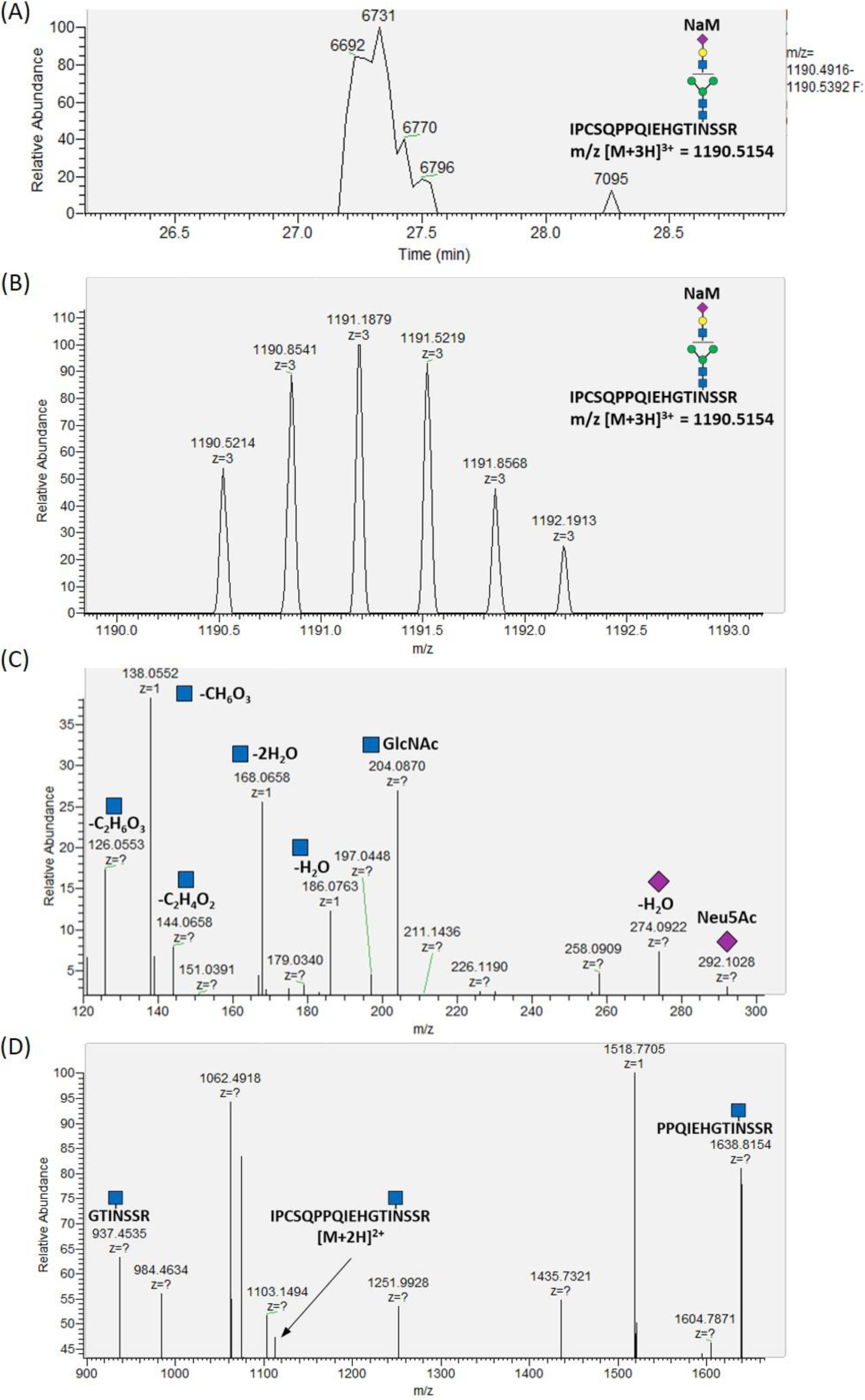
Mass-spectrometric verification of the identified NaM-glycosylation of the IPCSQPPQIEHGTINSSR glycopeptide on MS^1^ **(A+B)** and MS^2^ level **(C+D). (A)** Extracted ion chromatogram of the expected [M+3H]^3+^ m/z-ratio of 1090.5154 of the tryptic MNa-IPCSQPPQIEHGTINSSR glycopeptide on MS^1^ level. **(B)** Verification of the charge state (z=3) of the detected glycopeptide via investigation from the isotope pattern. **(C)** MS^2^-based verification of *N*-glycan sialylation via the detection of *N-*acetylglucosamine (GlcNAc) and sialic acid (Neu5Ac) reporter ions with the following m/z-values: [GlcNAc]^+^ = 204.087, [GlcNAc - H_2_O]^+^ = 186.076, [GlcNAc - 2H_2_O]^+^ = 168.066, [GlcNAc - C_2_H_4_O_2_]^+^ = 144.065, [GlcNAc - CH_6_O_3_]^+^ = 138.055, [GlcNAc - C_2_H_6_O_3_]^+^ = 126.055), [Neu5Ac]^+^ = 292.103 and [Neu5Ac - H_2_O]^+^ = 274.092. **(D)** Fragment ions of the glycopeptide. Blue square: *N-*acetylglucosamine (Gn, GlcNAc), green circle: mannose (M), yellow circle: galactose (A), purple rhombus: sialic acid (Na, Neu5Ac), Hex: hexose (yellow and green circle: stands for the presence of either mannose or galactose).

## Notes

https://dataview.ncbi.nlm.nih.gov/object/PRJNA665456?reviewer=olkq2ha7mnb640euv2rj04vqqo

